# Thiosulfinate tolerance gene clusters are common features of *Burkholderia* onion pathogens

**DOI:** 10.1101/2024.01.24.577064

**Authors:** Sujan Paudel, Mei Zhao, Shaun P. Stice, Bhabesh Dutta, Brian H. Kvitko

## Abstract

*Burkholderia gladioli* pv. *alliicola, B.cepacia*, and *B. orbicola* are common bacterial pathogens of onion. Onions produce organosulfur thiosulfinate defensive compounds after cellular decompartmentalization. Using whole genome sequencing and *in silico* analysis, we identified putative thiosulfinate tolerance gene (TTG) clusters in multiple onion-associated *Burkholderia* species similar to those characterized in other *Allium*-associated bacterial endophytes and pathogens. Sequence analysis revealed the presence of three *Burkholderia* TTG cluster types with both Type A and Type B being broadly distributed in *B. gladioli*, *B. cepacia*, and *B. orbicola* in both the chromosome and plasmids. Based on isolate natural variation and generation of isogenic strains, we determined the *in vitro* and *in vivo* contribution of TTG clusters in *B. gladioli*, *B. cepacia*, and *B. orbicola*. The *Burkholderia* TTG clusters contributed to enhanced allicin tolerance and improved growth in filtered onion extract by all three species. TTG clusters also made clear contributions to *B gladioli* foliar necrosis symptoms and bacterial populations. Surprisingly, the TTG cluster did not contribute to bacterial populations in onion bulb scales by these three species. Based on our findings, we hypothesize onion-associated *Burkholderia* may evade or inhibit the production of thiosulfinates in onion bulb tissues.

## Introduction

Onion (*Allium cepa* L.) production industry, valued at US $1 billion is a major contributor to the USA economy (Belo et al., 2023). Bacterial pathogens can infect onions at different stages of production from seedling to storage posing a serious challenge to growers (Mark et al., 2002; Zhao et al., 2022). Upon conducive environmental conditions, bacterial pathogens can cause more than 50% losses in production (Belo et al., 2023). Members in the *Burkholderia* genus are historically associated with onion disease with its first report dating back to 1940s (Burkholder, 1942). *Burkholderia cepacia* complex (Bcc) group causes sour skin of onion whereas *Burkholderia gladioli* pv. *alliicola* (Bga) causes slippery skin (Burkholder, 1950). In addition to the common onion pathogenic members *B. cepacia, B. cenocepacia,* and *B. ambifaria*, three new onion pathogenic species *B. orbicola, B. semiarida,* and *B. sola* were recently described as members of the Bcc complex (Morales-Ruíz et al., 2022; Velez et al., 2023). While all described onion pathogenic members in *Burkholderia* infect bulbs and are storage pathogens, Bga is also associated with foliar necrosis (Kawamoto and Lorbeer, 1974; Lee et al., 2005).

Plants utilize different chemical compounds as a defense response against the attack of pathogens and pests. In many cases, the inactive form of these compounds is converted into the active form only after the pathogen/insect attack. Onion, garlic, and other *Allium* species produce rapid and potent organosulfur compounds as a consequence of tissue damage (Lancaster and Collin, 1981; Rose et al., 2005). In garlic, the thiosulfinate allicin produced by the reaction of alliinase on alliin is a primary volatile antimicrobial compound that inactivates enzymes and depletes the reduced glutathione pool. Allicin is the compound responsible for the odor of crushed garlic. Onion, on the other hand, when disrupted produces asymmetric 1-propenyl methyl thiosulfinates in onion extracts. The action of alliinase and another enzyme, lachrymatory factor synthase (LFS) converts isoalliin into syn-propanethial-S-oxide, which is an irritant that induces tears (Silvaroli et al., 2017).

A plasmid-borne cluster of eleven-genes named the allicin tolerance (*alt*) cluster was found to confer increased tolerance to allicin and enhance virulence in *P. ananatis* on onion (Stice et al., 2020). Mutation in the *alt*/TTG region resulted in a smaller clearing zone in red onion scale necrosis assay and caused approximately 100-fold reduced bacterial population in onion tissue, and increased sensitivity to allicin and endogenous onion thiosulfinates. The PNA 97-1 mutant strain lacking the *alt* TTG cluster was severely reduced in the bacterial colonization of onion bulbs and scales suggesting the role of *alt* clusters in the virulence of the bacterium. Multiple chromosomal TTG clusters conferring allicin tolerance were also characterized in the garlic saprophyte *Pseudomonas fluorescens* PfAR-1, with similar clusters identified in the garlic pathogen *Pseudomonas salomonii* (Borlinghaus et al., 2020; Stice et al., 2020). The *Pantoea* and *Pseudomonas* TTG clusters had similar gene content but dissimilar gene cluster syntenies.

TTG clusters have not been reported in onion pathogenic members of *Burkholderia* genus. The *Burkholderia* pathogens are likely exposed to thiosulfinates during the infection process. In this study, we sequenced representative onion-isolated *Burkholderia* strains from Georgia, USA., and explored the distribution and function of putative TTG-like clusters from *Burkholderia*. Using allicin Zone of Inhibition (ZOI) and onion juice growth assays, we demonstrated the contribution of the putative TTG-like cluster to allicin tolerance and filtered onion extract growth in multiple onion-associated *Burkholderia* species. Engineered TTG mutant and TTG encoding heterologous expression plasmid derivatives were generated to determine the contribution of TTG cluster to onion foliar/red scale necrosis and *in planta* bacterial populations. Our results suggest that the TTG cluster contributes to variable virulence roles in onion depending on onion tissue-type and *Burkholderia* species.

## Results

### Identification of putative TTG-like cluster in *Burkholderia* species

Using *Pantoea ananatis* (Pan) PNA 97-1R TTG cluster gene sequences as a query, we performed a multigene blastX analysis to check the presence or absence of corresponding protein homologs in *Burkholderia*. Hits were obtained in the *Burkholderia* genus for multiple TTG gene sequences in the reference Pan PNA 97-1R *alt* TTG cluster. Further analysis of the hits in the whole genome sequence of *B. cepacia* 561 revealed the presence of putative TTG genes *altB*, *altC*, *altA*, *altE*, *altR*, *altI*, and *altJ*. The *altA, altB,* and *altC* genes are predicted to function as putative thiol/oxidoreductases, and *altJ* and *altE* encode for predicted peroxidase-like enzymes. The *altI* gene encodes for a putative carbon-sulfur lyase and *altR* encodes a putative repressor. Homolog of Pan TTG genes *altD, altG*, *altH, altJ,* and *gorB* were not found in the screened *B. cepacia* cluster. Similar TTG cluster genes were identified in the onion-isolated Bga strain 20GA0385. Clinker gene synteny analysis revealed the seven TTG genes were conserved in both Bga strain 20GA0385 and Pan strain PNA 97-1R. Amino acid (aa) percent identity between the two strains for *altB, altA,* and *altC* genes was 81%, 67%, and 37%, respectively. The peroxidase-like *altE* and *altJ* shared 63% and 29% aa identities, respectively. The putative C-S lyase encoding *altI* and the putative repressor *altR* shared 48% and 39% identity, respectively. The gene synteny between the two clusters; however, was not conserved (Figure 1).

**Figure 1:**
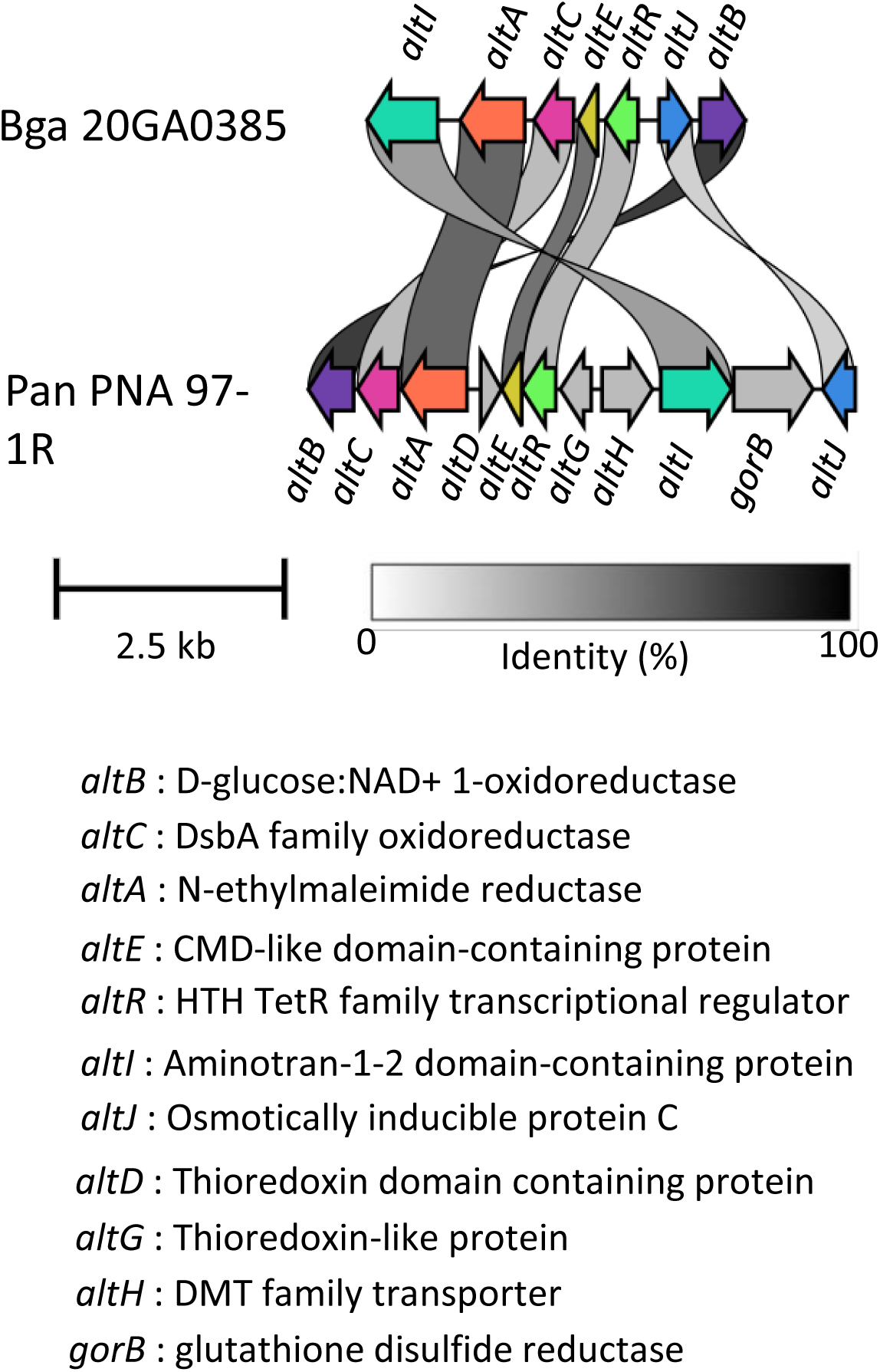
Gene cluster organization representing seven genes in the putative Thiosulfinate Tolerance Gene (TTG) cluster in *Burkholderia gladioli* pv. *alliicola* strain 20GA0385. Gene synteny of the Bga cluster relative to *Pantoea ananatis* strain PNA 97-1 is shown using the Clinker arrow diagram. Genes are color-coded based on nucleotide homology between the two gene clusters, the percentage of which is shown in the gradient scale. The proposed function of the genes in the cluster based on Stice et al., 2020 is listed.

### TTG clusters are widespread in *Burkholderia* species commonly isolated from onion

To study the distribution of putative TTG clusters in onion-associated strains, we conducted whole genome sequencing and assembly of 66 *Burkholderia* strains isolated from symptomatic onion. The species of sequenced strain was confirmed using the Type Strain Genome Server platform (Meier-Kolthoff and Göker, 2019). Out of the 66 sequenced strains, 22 were identified as *B. orbicola*, 20 were *B. cepacia,* 20 were *B. gladioli,* and 4 were *B. ambifaria* (Supplementary Table S3). Putative TTG-like clusters were found in 55 of the sequenced strains. All sequenced *B. orbicola* and *B. ambifaria* strains had TTG clusters. Among the 11 TTG negative strains, 2 were from *B. cepacia* species and 9 were from *B. gladioli* species. Interestingly, we discovered four strains that had two putative TTG clusters in a single strain, with three of these strains belonging to *B. orbicola* and one to *B. gladioli* (strain 20GA0350) (Figure 2).

**Figure 2:**
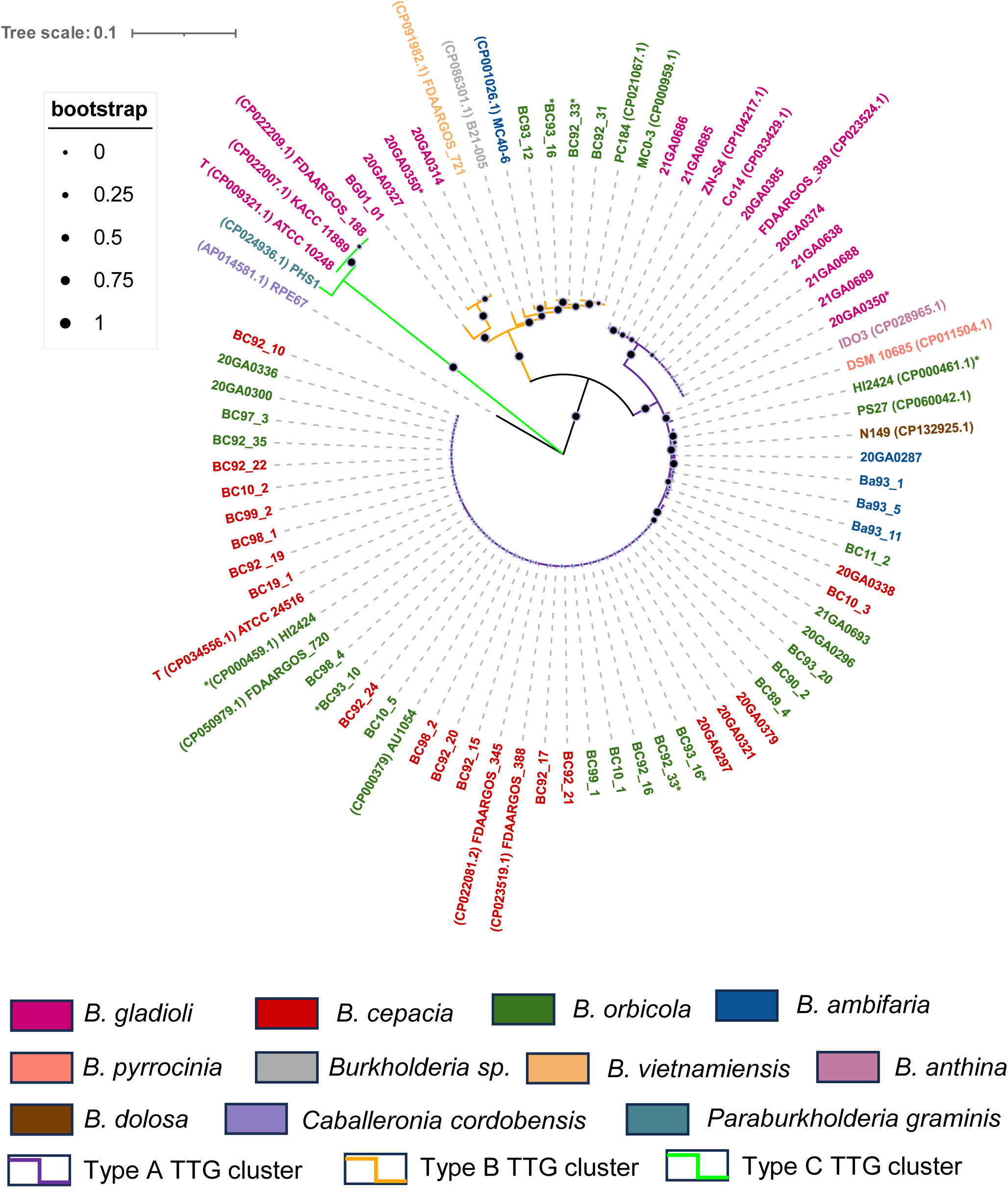
TTG cluster is widely distributed in *Burkholderia* species. A maximum-Likelihood method-based phylogenetic tree of n= 4952-bp partial TTG cluster consensus nucleotide sequences corresponding to the *altA*-*altB* gene region derived from whole genome sequenced *Burkholderia* isolates in the study and NCBI GenBank extracted TTG cluster sequences. 1000 bootstrap replicates were used as a test of phylogeny. Bootstrap support value on the scale of 0 to 1 is represented as a circle on the branch. Strain labels are color-coded based on species and branches are color-coded based on TTG cluster type. The accession number of the GenBank extracted sequences is provided in parentheses next to the strain name. Strains with two putative TTG clusters is noted with an asterisk next to its name. Letter T indicates a Type strain. The tree is rooted in the *Caballeronia cordobensis* strain RPE67 branch. Strain 20GA0329, 20GA0341, and TTG cluster Type B sequence of *B. gladioli* strain BC93_10 are omitted from the alignment as the entire TTG sequence was not present in a single contig. Information about the strains used is presented in Supplementary Table S3. Identity confirmation of the strains extracted from the NCBI GenBank database was confirmed using the TYGS webserver.

### Three distinct TTG cluster types are found based on nucleotide homology

To analyze the distribution and phylogeny of the TTG clusters, the identified putative TTG clusters were extracted from the respective genomes and aligned against each other. An additional 24 *Burkholderia* TTG sequences from closed genomes in the NCBI GenBank database were included in the alignment. The synteny of TTG cluster genes *altI* to *altB* was conserved for all the sequenced strains in our study. Among the NCBI GenBank extracted sequences, a slight difference in gene arrangement was observed in three *B. gladioli* strains and one *Paraburkholderia graminis* strain. The putative C-S lyase gene *altI* was present downstream of *altB* gene in the four strains. In the rest of the strains, the *altI* gene was present upstream of *altA* gene. The maximum-likelihood phylogenetic tree revealed the presence of three distinct TTG clades among the strains (Figure 2). The first branch had 63 TTG sequences with relatively less diversity among them. The second cluster had 13 sequences and the third cluster had four TTG sequences from *B. gladioli* and *P. graminis* species. The three clusters were named Type A, Type B, and Type C TTG clusters based on nucleotide homology and branching in the phylogenetic tree. The *B. gladioli* strains in the Type A and Type B branches formed a distinct sub-clade and showed species-specific branching (Figure 2). No species-specific branching was observed for other *Burkholderia* species. Among strains with two putative TTG-like clusters, three *B. orbicola* strains had one TTG cluster sequence each in Type A and Type B branches while *B. orbicola* strain HI2424 had both clusters in the Type A clade.

### The TTG-like cluster in *Burkholderia* contributes to allicin tolerance *in vitro*

To test the functional role of TTG-like clusters in sequenced strains, an allicin zone of inhibition (ZOI) assay was performed. Firstly, representative natural variant strains in *B. gladioli*, *B. cepacia*, and *B. orbicola* species with or without endogenous TTG clusters were tested for allicin tolerance. The TTG negative variant in all three species was highly sensitive to allicin based on the larger relative zones of inhibitions compared with the strains possessing the endogenous TTG cluster in the respective species group. (Figure 3A, 3D, 3F). There was no significant difference in ZOI area between endogenous Type A and Type B cluster types in *B. gladioli* and *B. orbicola* representative strains (Figure 3A, 3F). The presence of two TTG clusters in a single strain was not observed to contribute to significantly higher allicin tolerance in *B. gladioli* and *B. orbicola* (Fig 3A, D, F). The functional role of the TTG cluster was further assessed by engineering a TTG deletion mutant in the Bga strain 20GA0385. The TTG mutant was more sensitive to allicin than the WT strain (Figure 3B). Although extensive attempts were made, we were unable to generate corresponding TTG mutants in *B. cepacia* and *B. orbicola* in part due to their high intrinsic resistance to aminoglycoside antibiotics creating challenges for clean selection and recalcitrance to *sacB*-mediated counter-selection. Thus we cloned the Type A TTG cluster (from *B. orbicola* strain 20GA0385) and TTG Type B cluster (from *B. orbicola* strain BC93_12) into pBBR1MCS-2 for complementation and heterologous expression in strains lacking a native TTG cluster. Plasmid-based expression of either cluster type in the TTG mutant background restored the ZOI phenotype to WT level. The TTG Type B complementing plasmid conferred significantly higher allicin tolerance compared to the WT strain and the TTG Type A complementation clones (Fig 3B). To test if the functional role of TTG cluster is conserved across major onion pathogenic *Burkholderia* species, we transformed TTG Type A and Type B expression plasmids separately into TTG negative natural variant backgrounds: BC83-1 (*B. cepacia*), LMG 30279^T^ (*B. orbicola*), and BG92_3 (*B. gladioli*). The TTG expression isogenic lines were significantly more tolerant to allicin in all three species groups compared to their respective TTG negative WT (Fig 3C, E, G). In *B. cepacia* BC83_1, TTG Type B expression plasmid contributed significantly to enhanced allicin tolerance as compared to the TTG Type A (data not shown) and the WT strain (Figure 3E). No statistical difference in allicin tolerance was observed between Type A and TTG Type B expression plasmids in *B. orbicola* strain LMG 30279 and *B. gladioli* strain BG92_3 (Figure 3C, 3G).

**Figure 3:**
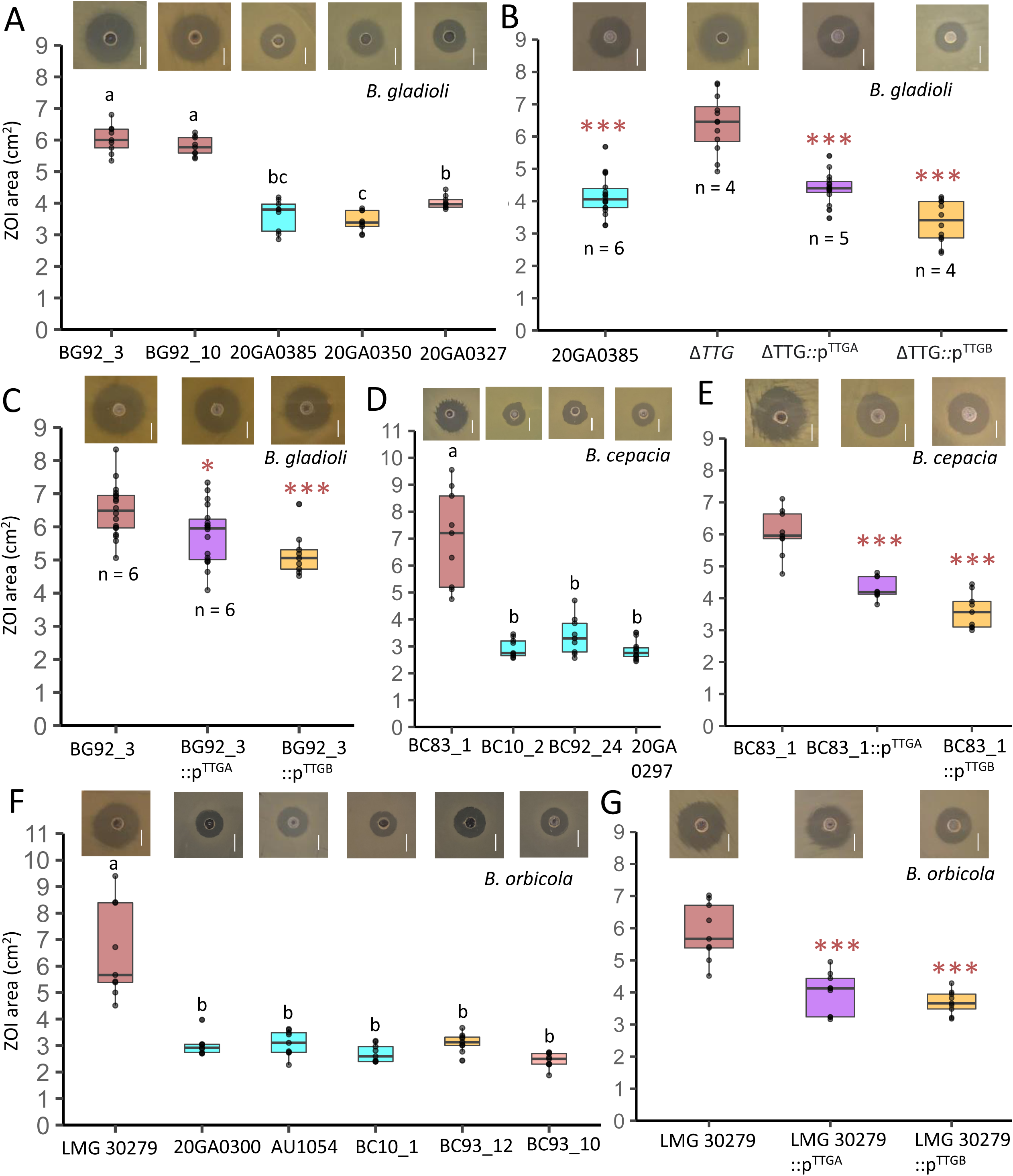
TTG cluster contributes to allicin thiosulfinate tolerance in *Burkholderia* species. **A**, Allicin zone of inhibition (ZOI) area of select natural variant isolates of *B. gladioli;* **B** ZOI area of engineered TTG mutant and Type A and TTG Type B cluster complementing strains of *B. gladioli* strain 20GA0385. **C,** ZOI area of TTG negative *B. gladioli* strain BG92_3 relative to its TTG Type A and Type B plasmid derivatives. **D** ZOI area of select natural variant isolates of *B. cepacia.* **E,** ZOI inhibition area of TTG negative *B. cepacia* strain BC83_1 compared to its TTG Type A and Type B plasmid derivatives. **F,** ZOI inhibition area comparison of representative natural variants of *B. orbicola.* **G,** ZOI inhibition area of *B. orbicola* TTG negative type strain LMG 30279 and its Type A and TTG Type B plasmid derivatives. Cyan colored bar represents a strain with endogenous TTG Type A cluster, brown bar represents a TTG negative engineered or natural variant strain, the purple bar represents TTG negative strains with TTG Type A expression plasmid, the orange bar represents TTG negative strain with TTG Type B expression plasmid or strains with endogenous TTG Type B cluster and salmon bar represents strain with both Type A and TTG Type B cluster. Representative images of ZOI plates tested with corresponding treatments are presented above the box. The scale bar in the image represents 1 cm length. Significance grouping based on ANOVA followed by Tukey’s post-test is shown as letters above the boxes in natural variants experiments. A pairwise t-test was done to determine the significant differences in ZOI area between the isogenic TTG-lacking strains and plasmid-expressed TTG cluster strains. Level of significance: 0 ‘***’ 0.001 ‘**’ 0.01 ‘*’ 0.05. The experiment was repeated at least three times or as represented by n = number of independent experimental repeats. The depicted data point represents all biological replicates from independent experimental repeats. Each experiment was conducted with three biological replicates.

### TTG-negative strains have impaired growth in filtered onion juice

Stice et al. 2020 demonstrated that endogenous onion thiosulfinates in onion juice restricted bacterial growth in a manner similar to synthesized allicin. To test if endogenous thiosulfinates affect the growth of TTG derivatives in different *Burkholderia species*, we conducted a filtered onion juice growth assay. In half-strength onion juice diluted with water, the Bga 20GA0385 TTG mutant had a dramatic growth slower than the WT strain over 48 h. The growth of the TTG mutant improved significantly when complemented with the plasmid harboring TTG gene cluster (Figure 4A). The TTG Type A and Type B plasmid derivatives grew significantly better in onion juice than the TTG negative natural WT variant in *B. cepacia* and *B. orbicola* (Fig 4B, 4C). When analyzed across the three experimental repeats, the OD_600_ values recovered for *B. orbicola* LMG 30279 Type B strain were significantly higher compared to the TTG Type A strain derivative 16 to 48 hours post-inoculation (Supplementary Table S4). The *B. cepacia* TTG Type B derivative growth was similar to the TTG Type A strain over 48 h across the three experimental repeats (Supplementary Table S4).

**Figure 4:**
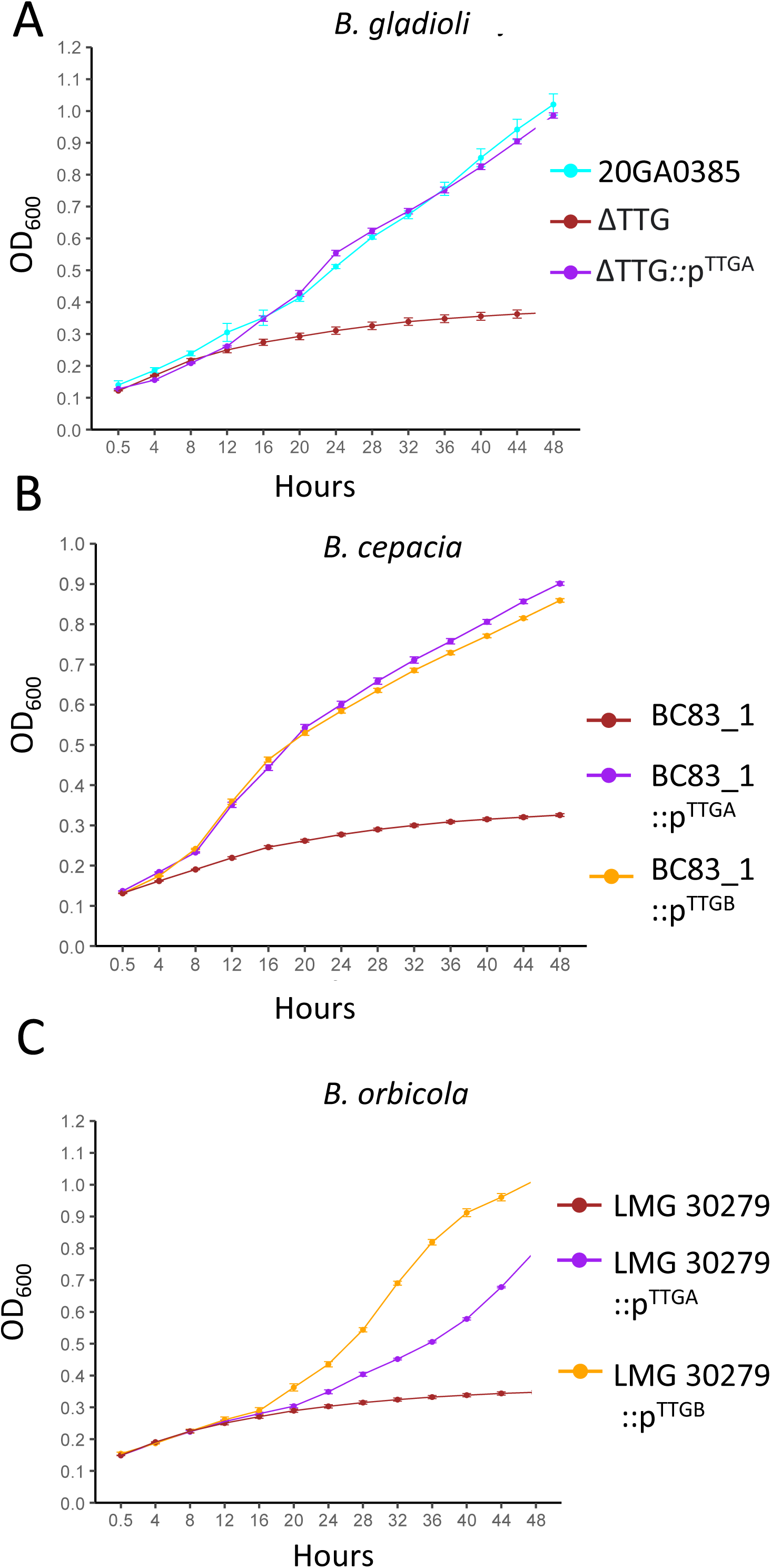
TTG negative strains are impaired in onion juice growth. Growth comparison of **A,** *B. gladioli* strain 20GA0385, TTG mutant derivative and TTG Type A complementing strain **B,** *B. cepacia* TTG negative natural variant BC83_1 and TTG Type A and Type B plasmid derivatives **C,** *B. orbicola* natural variant type strain LMG 30279 and TTG Type A and Type B plasmid derivatives, *in* half strength yellow onion juice for 48 h. Error bars represent ± Standard Error. Representative experimental data out of three performed independent experimental repeats are shown in the figure. Color coding is as in Figure 3. An average of six technical replicates per treatment is shown per time point. The Y axis crosses at 0.5 h post-inoculation. The test of significance was done with recorded OD_600_ values at each time point in between the treatment strains and is presented in Supplementary Table S4.

### TTG cluster in *B. gladioli* pv. *alliicola* contributes to foliar necrosis and bacterial populations

The *B. gladioli* TTG mutant and the TTG plasmid derivatives in TTG-negative WT background were tested for their contribution to onion foliar necrosis and bacterial populations. The 20GA0385 ΔTTG was significantly reduced in onion seedling lesion length and bacterial population compared to the WT strain. The necrosis length and bacterial population were restored to the WT level when the TTG mutant was expressed with a TTG Type A complementing plasmid (Fig 5A, 5B). The TTG Type A plasmid also contributed to significantly higher necrosis length and bacterial populations when transformed in a TTG-negative natural variant strain BG92_3 (Fig 5A, 5B). In addition, a time course bacterial population growth assay for both Bga 20GA0385 and ΔTTG was performed. Bacterial populations at six hours post-inoculation were in the range of 10^4^ CFU/mg of onion tissue. Bacterial populations for the WT strain increased gradually and reached the maximum plateau from day 3 to day 5 post-inoculation in the range of 10^6^-10^8^ CFU per mg of infected leaf tissue. The bacterial population recovered for ΔTTG Day 3 to Day 5 post-infection was significantly lower compared to the WT strain (Fig 6A).

**Figure 5:**
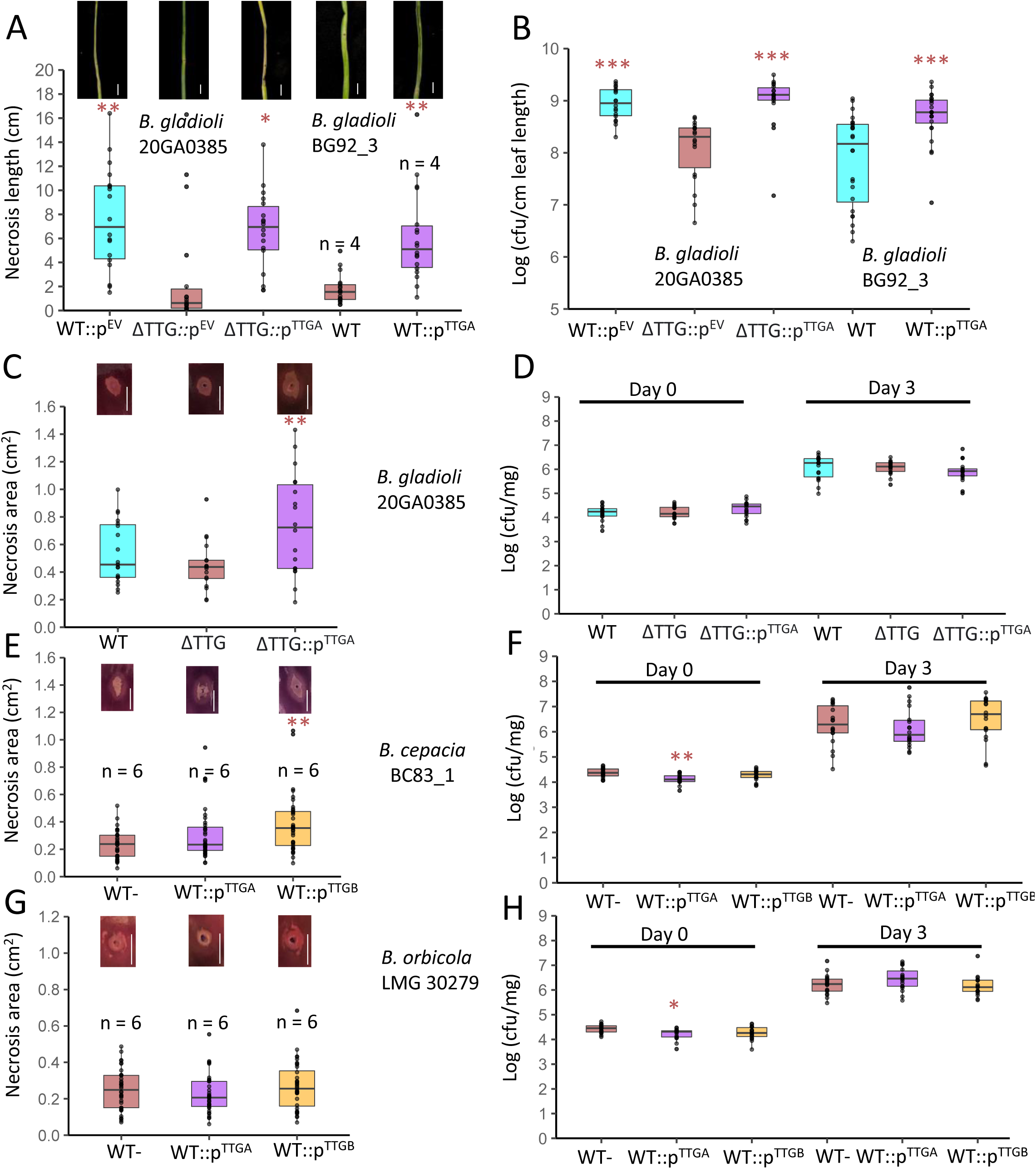
Onion foliar /Red Scale Necrosis (RSN) assay and corresponding *in planta* population quantification of natural variants, engineered mutants, and TTG plasmid transformed strains of three *Burkholderia* species. **A, B** Box plot showing onion foliar necrosis length and *in planta B. gladioli* bacterial population in onion leaf tissue at 3 days post inoculation (dpi). EV = Strain carrying empty vector pBBR1MCS-2. Box plot showing RSN area and *in planta* bacterial population count log (CFU/mg) of onion scale tissue at 4 h and 3 dpi for representative samples **C, D** *B. gladioli* strain 20GA0385, its engineered TTG mutant derivative and TTG Type A complementation clone; **E, F** *B. cepacia* TTG negative natural variant BC83_1 and TTG Type A and Type B plasmid derivatives; **G, H** *B. orbicola* TTG negative natural variant Type Strain LMG 30279 and TTG Type A and Type B plasmid derivatives. Color coding is described in Figure 3. The significant difference in necrosis length, RSN necrosis area, and in planta bacterial load of endogenous TTG cluster or plasmid-based TTG cluster harboring strain was compared to its engineered TTG mutant or TTG negative Wild Type (WT) natural variant using pairwise t-test applying Bonferroni coefficient. Each jitter point represents a biological replicate of all the experiments. The image above the box plot highlights tissue necrosis from representative samples for corresponding inoculated treatment. The scale bar in the figure represents a 1 cm length. The experiment was repeated at least three times or as represented by n = number of independent experimental repeats. Each experiment had six technical replicates. Level of significance: 0 ‘***’ 0.001 ‘**’ 0.01 ‘*’ 0.05

**Figure 6:**
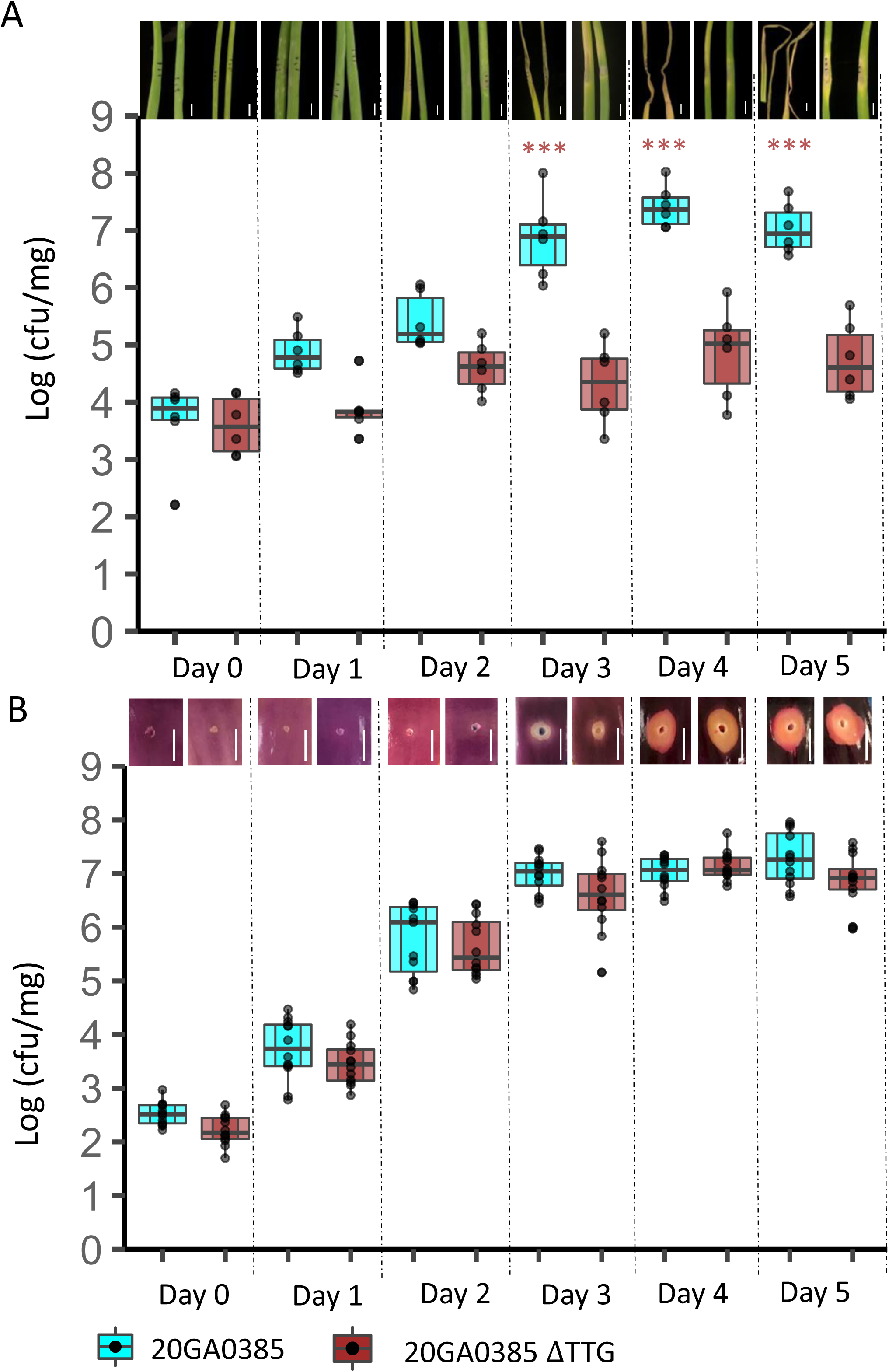
*B. gladioli* TTG cluster contributes to onion foliar bacterial load. In planta population quantification of TTG mutant and WT 20GA0385 strain from **A,** onion leaves and **B,** Red scales from Day 0 (5 hr) to Day 5 post-infection. A representative image of leaves or scales inoculated with ∼2.4 x 10^6^ CFU of WT and TTG mutant strain at each dpi is shown above the box plot. Significant differences in log (CFU/mg) of leaf or scale tissue between the two treatments for each day of sampling were calculated using a pairwise t-test applying the Bonferroni probability adjustment method. Level of significance: 0 ‘***’ 0.001 ‘**’ 0.01 ‘*’ 0.05. The scale bar represents 1 cm in length. The experiment was repeated two times.

### TTG cluster in *Burkholderia* does not contribute to symptom production or bacterial populations in onion scale tissue

The Red Scale Necrosis (RSN) assay was conducted with strains in three *Burkholderia* species to study the role of TTG cluster in lesion area and scale bacterial population. The *B. gladioli* 20GA0385 ΔTTG was not altered in the RSN area compared to the WT. The TTG Type A complementing plasmid clone had a significantly larger necrosis area compared to the TTG mutant. Bacterial populations for three *B. gladioli* 20GA0385 treatments (WT, ΔTTG, and ΔTTG::p^TTGA^) were not significantly different from each other 4 h and 3 days post-inoculation (Fig 5D). In the case of the *B. cepacia* strain BC83_1, the TTG Type B expression clone contributed to the RSN area, but the TTG Type A clone did not affect the RSN area (Fig 5E). Onion scale bacterial population for *B. cepacia* and *B. orbicola* TTG Type A expression clone was significantly reduced at 4 hours post-inoculation compared to the respective WT and TTG Type B expression clones (Fig 5F, 5H). No significant difference in bacterial population and the RSN area was observed for *B. cepacia* and *B. orbicola* WT and TTG plasmid derivatives at day 3 post-infection (Fig 5F, 5G,5H). Onion scale bacterial population for *B. gladioli* 20GA0385 WT and ΔTTG strains from day 0 to day 5 were not significantly different from each other (Figure 6B).

## Discussion

Using whole genome sequencing and phenotypic assays, we studied the distribution and functional role of TTG-like clusters in *Burkholderia* species commonly isolated from onion. RSN and foliar necrosis assays were conducted to investigate the role of TTG-like clusters in tissue-specific symptom production and bacterial population. A putative TTG-like cluster was identified in onion-isolated Bga strain 20GA0385. The genes present in the cluster had similar predicted functions compared to TTG cluster genes in *P. ananatis* strain PNA 97-1 but the gene synteny and orientation were different (Figure 1). Whole genome sequencing revealed the widespread distribution of TTG-like clusters in onion-isolated *Burkholderia* species. Maximum likelihood-based phylogeny of whole genome sequenced and NCBI GenBank extracted representative TTG sequences highlighted the presence of three distinct clades (Figure 2). These three clades were named Type A, Type B, and Type C TTG clusters based on nucleotide homology. The TTG Type A and Type B clusters both contributed to allicin tolerance in tested *B. gladioli*, *B. cepacia*, and *B. orbicola* strains (Figure 3). The clusters also contributed to improved bacterial growth in half-strength onion-filtered extract (Figure 4). In the onion foliar assay, the Bga 20GA0385 TTG cluster contributed to necrosis length and bacterial *in planta* population count (Figure 5A, 6A). The TTG Type A cluster when expressed in the *B. gladioli* TTG negative strain BG92_3 background contributed significantly to onion foliar necrosis length and *in planta* bacterial population count (Figure 5A). The TTG clusters Type A and Type B, however, were not required for red-scale necrosis area and scale bacterial load in *B. gladioli* and *B. orbicola* (Figure 5). The TTG Type B cluster in *B. cepacia* strain BC83_1 contributed significantly to the RSN area but was inconsequential for the red scale *in planta* population count. The *B. cepacia* BC83_1 TTG Type A cluster was not required for the RSN area or in-planta bacterial load (Figure 6E,6F).

The role of the TTG cluster in conferring allicin tolerance in bacterial onion pathogen *P. ananatis* is well correlated with the bacterium’s ability to colonize necrotized onion tissue. As members of the bacterial genus *Burkholderia* are routinely isolated from infected onion, we hypothesized they might utilize a similar strategy to combat and colonize the hostile onion environment. We sequenced and assembled the genomes for a panel of onion-isolated *Burkholderia* strains collected over 40 years from Georgia, USA. A majority of the sequenced strains belonged to the three common onion-associated *Burkholderia* species: *B. gladioli*, *B. cepacia*, and *B. orbicola*. Putative TTG-like clusters were distributed in all three species. No congruence was observed between the core-gene-based species tree and the TTG cluster-based tree suggesting the TTG clusters might be horizontally acquired. Multiple genes in *P. ananatis* TTG cluster have also been predicted to be involved in horizontal gene transfer (Stice et al., 2018). Upon alignment of the TTG sequence region, three distinct groups were formed based on homology with the consensus. The grouping was also reflected in the phylogenetic tree. The Type A cluster was the most common while no Type C cluster was found in our sequenced panel of strains. Similarly, no Type B cluster was found in *B. cepacia* strains. Four strains harbored two putative TTG clusters simultaneously in their genome. The presence of multiple TTG clusters in the same strain is not exclusive to *Burkholderia* genus. The *Pseudomonas fluorescens* strain *Pf*AR-1 has three copies of TTG clusters that confer resistance to allicin when expressed heterologously in *E. coli* (Borlinghaus et al., 2020). The *B. orbicola* strain HI2424 harbors two Type A TTG clusters, one on the chromosome and the other in the plasmid. The Type C cluster was found in four strains deposited in the NCBI GenBank database but was not identified in our strain panel. The three strains harboring Type C cluster were from *B. gladioli* and they grouped closely to TTG cluster from *Paraburkholderia graminis* strain PHS1. The number of nucleotide substitutions in the Type C TTG strains suggests the TTG cluster might have existed before the differentiation of *Paraburkholderia* genus from *Burkholderia.* The Type C cluster type was peculiar in the sense that the putative *altI* C-S lyase encoding gene was present downstream of *altJ* gene and transcribed in the same direction as *altJ*. Apart from this gene, the gene organization of other genes in the *Burkholderia* TTG cluster is conserved. One clinical *B. gladioli* strain BCC0507 had an insertion of a putative IS3 family transposase CDS in between the *altA* and *altC* gene (data not shown).

The gene synteny of the TTG cluster among different bacterial genera is not conserved. The *Burkholderia* TTG cluster is compact with seven genes as opposed to the bacterial onion pathogen *P. ananatis* which has eleven genes in its TTG cluster. The *gor* gene encoding glutathione reductase is associated with TTG clusters in some *Pantoea, Erwinia*, and *Pseudomonas* species but is absent in the *Burkholderia* TTG cluster (Couto et al., 2016; Borlinghaus et al., 2020). Similarly, three additional TTG genes *altD, altH*, and *altG* present in *P. ananatis* PNA 97-1 are absent in *Burkholderia.* The TTG cluster in *P. ananatis* shares some notable differences in gene synteny with the TTG cluster described in representative strains from *Pseudomonas* genus. Despite the differences between the two genera, the TTG cluster gene synteny is conserved among the *P. fluorescens* strain *Pf*AR-1, *P. salomonii* strain ICMP 14252, and *P. syringae* pv. *tomato* strain DC3000 within the *Pseudomonas* genus.

Although widely distributed in onion-isolated bacterial species, it is important to note that the TTG cluster is not exclusive to onion-pathogenic species. Members in the *P. fluorescens* group are common saprophytes. Analysis of the TTG clusters in *Burkholderia* species deposited in NCBI GenBank database suggests the putative cluster is present in strains isolated from diverse ecosystems such as reactor sludge, clinical patient, soil, sputum, and plant hosts such as *Dendrobium* and *Gladiolus.* As the sequenced strains in our study are draft genomes, we were not able to definitively determine the chromosomal or plasmid origin of the TTG cluster. We used TTG clusters from NCBI closed genomes as a reference to analyze the genomic context of TTG clusters in our panel. In all *B. gladioli* and *B. cepacia* closed genomes in the database, the TTG cluster was harbored in a plasmid. *B. orbicola* strains harbored TTG clusters in both chromosomes or plasmids. We used BLAST analysis to compare 35 kb regions in the *B. cepacia* ATCC 25416 including the TTG Type A clusters, and its upstream and downstream gene region against the whole genome sequenced strains in our panel. The gene synteny of the 35 kb region was conserved and shared high nucleotide identity among the 24 strains harboring TTG clusters. Out of the 24 hits obtained from BLAST analysis, 6 were from *B. cepacia* strains and the remaining 18 were from *B. orbicola* species. The 35 kb plasmid region harbored in *B. cepacia* strain ATCC 25416 was also 100% identical to the region harbored in chromosome 2 of *B. orbicola* strain HI2424. Although no conclusion can be drawn on the origin of TTG clusters in these strains, it can be predicted that the TTG clusters can be present in both chromosome or plasmids in *Burkholderia* strains.

The presence of two TTG cluster types in a single *Burkholderia* strain was found in four instances. When two of these strains were tested against the natural variant strains harboring a single or no TTG cluster using the allicin ZOI assay, enhanced resistance to allicin was not observed. This was in contrast to *P. fluorescens* strain *Pf*AR-1 and garlic pathogen *P. salomonii* strain ICMP14252, where the presence of two or more endogenous TTG clusters correlated with enhanced tolerance compared to the single TTG cluster harboring *P. syringae* pv. *tomato* strain DC3000. The TTG Type B cluster conferred enhanced tolerance to allicin in *B. gladioli* strain 20GA0385 and *B. cepacia* strain BC83_1 compared to Type A TTG and WT strains. The enhanced phenotype by Type B cluster was not observed in *B. orbicola* strain LMG 30279 and *B. gladioli* strain BG92_3. Although variation was seen among the experimental repeats, we saw a qualitative growth difference between *B. cepacia* TTG Type A and Type B clusters in a couple of onion-filtered extract growth assay repeats (Figure 4B). The *B. orbicola* TTG Type B cluster grew significantly better in onion extract compared to the TTG Type A cluster (Supplementary Table S4). It is not clear what might have contributed to enhanced tolerance by TTG Type B clusters as gene synteny is conserved in both cluster types.

The virulence role of the TTG cluster was analyzed using onion foliar and red scale necrosis and population assays. We observed contrasting virulence roles of *B. gladioli* TTG cluster depending on onion tissue type. The *in planta* foliar population count for Bga 20GA0385 WT strain increased consistently from Day 0 and reached the plateau on Day 3 (Figure 6A). The foliar necrosis symptoms by Bga WT strain started appearing on Day 2 and by Day 3, the necrosis extended across the tip with leaves appearing wilted and turned brittle in the subsequent days. The foliar population levels for the TTG mutant remained constant throughout the sampled period. The necrosis symptoms observed with the TTG mutant were restricted around the region of the point of inoculation. Although a 100-fold diluted suspension was used for scale time course assay, the population count for both WT and TTG mutant in red scale reached the peak at the same time as in the foliar tissue (Figure 6B). The necrosis area for the majority of both WT and TTG inoculated scales continued to increase even after the population count reached the plateau. Unlike in the foliar tissue, the TTG mutant population count from the red scale assay was comparable to WT across all the sampling points (Figure 6B). These findings from the *in planta* population assay, contradict the results of the onion filtered extract growth assay, where the TTG mutant was severely impaired in the onion juice growth compared to the WT strain (Figure 4A). The differences in *in vitro* and *in planta* experiment results suggest that bacteria in onion scale tissue may utilize mechanisms to inhibit the onion-produced thiosulfinates in necrotized tissue. Alternatively, as thiosulfinates are produced by mixing of CSO precursors and alliinase following the disruption of the vacuole, bacteria may exhibit a hemibiotrophic lifestyle keeping the cells intact in the early stages of invasion This is in line with what we observed in the red scale time course assay where the maximum necrosis area followed after the bacterial population reached the plateau. Based on these observations, we can speculate that the *Burkholderia* strain might be in a hemibiotrophic survival mode in onion scales.

The TTG clusters Type A and Type B were inconsequential for the population on red scale in both *B. orbicola* and *B. cepacia.* As we inoculated a relatively high number of bacteria (∼10^8^ CFU/ml) in the scales for all three tested species, we sampled a set of scales four hours post-inoculation to check if bacteria have enough threshold for growth inside the scale tissue. The bacterial count four hours post-inoculation was similar for all the treatments tested across the three species. Although not dramatic, we did see a significant population count reduction for TTG Type A plasmid derivative in both *B. cepacia* and *B. orbicola* compared to their respective WT strain. Surprisingly, this trend was not observed for the TTG Type B plasmid derivative that has the same plasmid backbone and was selected under the same conditions. The reduction in Day 0 bacterial population was not observed for *B. gladioli* treatments where a much lower antibiotic concentration (50 µg/ml) was used for selection. It is unclear why the Day 0 population reduction phenotype is specific only to TTG Type A plasmid derivative.

Through this study, we have shown the functional role of the TTG cluster is conserved in distantly related onion pathogenic members of the *Burkholderia* genus. Thiosulfinates are broad-acting antimicrobials. Bacteria in the *Burkholderia* genus may have acquired tolerance mechanisms to evade or inhibit the produced thiosulfinates in a tissue-specific manner.

Functional characterization of the TTG constituent genes and their interaction mechanisms with bacteria could open up details on how bacteria can survive and replicate in a challenging onion environment.

## Materials and Methods

### Bacterial growth conditions

Bacterial strains used for cloning, mutagenesis, and construction of the TTG expression plasmid in this study are listed in Supplementary Table S1. Primers and synthesized dsDNA fragments used in the study are listed in Supplementary Table S2. *Escherichia coli* strains DH5α and RHO5 and all *Burkholderia* strains used for the *in vitro* experiments were grown in LB (per liter, 10 g of tryptone, 5 g of yeast extract, 5 g of NaCl) broth or agar (15 g of agar) at 37°C and 28°C, respectively. Antibiotics and chemicals were supplemented with the growth media at the following final concentrations, per milliliter: 50 – 1000 µg of kanamycin, 10 µg of gentamicin, 100 – 200 µg of Diaminopimelic acid (DAP), 50 µg of X-Gal (5-bromo-4-chloro-3-indolyl-beta-D-galacto-pyranoside), 100 µg of Xgluc (5-bromo-4-chloro-3-indolyl-beta-D-glucuronic acid), and 40-60 µg of rifampicin, as appropriate.

### Plant growth conditions

Onion sets (*Allium cepa L.* cv. Century) were planted in 10 cm x 8 cm (diameter x height) plastic pots filled with SunGrow 3B potting soil and maintained at greenhouse conditions with 25–28°C, 12L:12D photoperiod for 5 months from January to May until inoculation. To grow the onion seedlings, onion seeds (*Allium cepa* var. Texas Grano 1015Y) were sown in (5 cm x 5 cm) pots with SunGrow 3B potting soil and maintained at the same conditions as described above in the greenhouse for 8-12 weeks.

### Identification of TTG-like cluster in *Burkholderia*

The nucleotide sequences of the individual genes in the TTG-like cluster in *P. ananatis* strain PNA 97-1R (NCBI Accession number: NZ_CP020945.2) were blasted against the *Burkholderia* genome database (taxid: 32008) in NCBI GenBank database using blastX web interface platform. The whole genome sequences of the hits obtained from the blastX search were screened for the presence of a putative TTG-like cluster. The locus tag numbers for the putative genes in the *Burkholderia* TTG-like alt cluster were determined using Geneious Prime v 2023.2.1.

The synteny analysis of the TTG cluster in Bga representative strain 20GA0385 was performed with Clinker web-based platform CAGECAT using *P. ananatis* PNA 97-1 (GenBank Accession: CP020945.2) TTG cluster as a reference (van den Belt et al., 2023). Bakta annotated TTG nucleotide clusters from PNA 97-1 and Bga strain 20GA0385 were used as input for CAGECAT synteny analysis (Schwengers et al., 2021). The identity threshold was set at 0.3 for the analysis.

### Whole genome sequencing of select *Burkholderia* strains

A total of 66 *Burkholderia* strains isolated from onions in different regions of Georgia state, USA were sent for whole genome sequencing using Novogene Co., Ltd (Beijing, China) and MicrobesNG Illumina sequencing (Microbes NG, Birmingham, UK). Raw sequences were filtered using fastp v 0.20.0 (Chen et al., 2018), and quality checks were conducted using fastqc v 0.11.9 (Andrews, 2010). The processed reads were assembled using SPAdes v 3.14 (--isolate --cov-cutoff auto mode) (Bankevich et al., 2012) and filtered for a minimum contig size of 500 bp. Assembled contigs were annotated using the Prokka annotation pipeline (Seemann, 2014). All whole genome sequences were uploaded to the NCBI GenBank database under BioProject ID PRJNA1048086. Strain metadata information is presented in Supplementary Table S3.

The species identity of the sequenced strains was confirmed using the Type Strain Genome Server (TYGS) (Meier-Kolthoff and Göker, 2019). Assemblies were uploaded to the server as inputs. A pairwise GGDC formula 2 (d_4_) value of >70% was used as the cutoff for species identification. Bga strain 20GA0385 was also confirmed using a 727 bp partial *recA* gene-based phylogeny (Supplementary Figure S1).

### Phylogeny of TTG*-*like clusters based on whole genome and NCBI extracted sequences

The presence/absence of TTG cluster sequence in the assembled genome sequences was determined using Map to Reference (Bowtie) and/or custom BLAST plugin in Geneious Prime v 2023.2.1 (Langmead, 2010). The TTG cluster sequence from *B. gladioli* pv. *alliicola* strain FDAARGOS_389 (GenBank: CP023524.1) was used as a query sequence for both Map to Reference and custom BLAST analysis. All genome sequenced strains were used to create a custom BLAST database and the query sequence was used as input against the custom database to perform the analysis. The matched regions obtained from the map to reference analysis and custom BLAST analysis were extracted and aligned using the MAFFT plugin in Geneious Prime. Putative *Burkholderia* TTG clusters from closed genomes in the NCBI GenBank database were also extracted and included in the alignment. Obtained hits smaller than 4952 bp corresponding to *altA* – *altR* gene region were excluded from further analysis. Metadata for the strains used in the analysis is presented in Supplementary Table S3. The aligned and trimmed sequence was used as input in MEGA X software for maximum likelihood method-based phylogenetic analysis (Kumar et al., 2018). Nucleotide substitution type with the Tajima-Nei substitution model was used and 1000 bootstrap replicates were used as a test of phylogeny. The obtained Newick (.nwk) tree was uploaded to the interactive Tree of Life (iTOL) database for further formatting (Letunic and Bork, 2021).

Identity confirmation of *B. gladioli* strains FDAARGOS_389 and 20GA0385 was done using a 727 bp partial *recA*-based phylogenetic analysis. Genome assemblies from representative strains in *B. gladioli* Clade 1A, 1B, 1C, 2, and 3 as described in (Jones et al., 2021) were downloaded from the NCBI GenBank database. Assemblies from *B. cepacia* strain ATCC 25416 and *B. cenocepacia* J2315 were also downloaded from the NCBI GenBank database. The presence of partial *recA* gene sequence was determined in the downloaded assemblies using Bowtie Map to Reference plugin in Geneious Prime v 2023.9.0, with the FDAARGOS_389 partial *recA* gene sequence as a query. The mapped regions in the target assemblies were extracted and aligned using the MAFFT multiple align plugin in Geneious Prime. The aligned sequences were used as input to generate a maximum likelihood based phylogenetic tree using MEGA-X. For the test of phylogeny, 1000 replicates of the bootstrap method were used. The identity of strains FDAARGOS_389 and 20GA0385 was confirmed by analyzing their grouping in the phylogenetic tree relative to reference strains from different clades as described in (Jones et al., 2021)

### Creation of *B. gladioli* TTG mutant strain

The unmarked deletion of the TTG cluster in Bga strain 20GA0385 was generated using an allelic exchange strategy. The 450 bp upstream (including 2 aa downstream of the stop codon in *altB* ORF) and 450 bp downstream flanking region (including 1 aa upstream of the stop codon in *altI* ORF) of TTG cluster from *B. gladioli* strain FDAARGOS_389 (GenBank: CP023522.1) along with attached attB1 and attB2 site was synthesized as a double-stranded DNA gblocks from Twist Biosciences. An AvrII restriction site was also included in between the deletion flank. The synthesized TTG fragment was BP cloned into suicide vector pR6KT2G using Gateway BP Clonase II enzyme mix (Thermo Fisher Scientific) (Stice et al., 2020). The cloned reaction following the treatment with Proteinase K was placed on the VMWP membrane (Millipore) and floated on top of sterile distilled water (dH_2_0) for 30 minutes. The de-salted mix was electroporated into *E. coli* MAH1 cells and transformants were selected on LB agar amended with gentamicin and Xgluc followed by incubation at 37^0^C overnight (Kvitko et al., 2012).

Selected transformants were grown overnight and the plasmid was prepped with GeneJet Plasmid Miniprep Kit (ThermoScientific, Watham, WA). The correct recombinant plasmid was confirmed with a HindIII restriction enzyme digest reaction and compared to a simulated pattern obtained using NEBcutter V2.0. One clone with the correct restriction digest size, was selected and sent for sequencing using pR6KT2GW-F and pR6KT2GW-R primer pair (Supplementary Table S2). The sequenced reads were aligned with the simulated vector created in Geneious Prime V 2021.1.6 to confirm the insert. The confirmed plasmid was then cloned into an LR-clonase compatible pK18mobsacB plasmid derivative pDEST1k18ms following the manufacturer’s recommendations (Mijatović et al., 2021). Following the Proteinase K treatment, the cloned reaction product was transformed into chemically competent *E. coli* DH5α cells, and the transformants were selected on LB amended with kanamycin, and screened for the correct insert using BsrGI restriction digest. The clone with the correct digest pattern was sent for sequencing with M13R49 primer and the sequence was analyzed as described above using Geneious Prime to confirm the insert. The confirmed insert was then transformed into E. competent *E. coli* RHO5 pir+ mating strain and selected on LB plate amended with DAP and kanamycin plates. *E. coli* no DAP liquid culture control and parental controls were also included. The plasmid with the deletion construct in the RHO5 strain was mated with WT 20GA0385 strain and merodiploids were selected on LB plate amended with rifampicin and kanamycin. For the sucrose-based counter selection, 10 merodiploid colonies were selected and suspended in 5 ml of sterile LB media. 50 µl of the suspension was spread on an LB plate amended with rifampicin and 10% 1M sucrose followed by incubation at 30^0^C for 48 hours. Isolated exconjugant colonies on the sucrose plate were patch-plated into both LB and LB amended with kanamycin plates to check the sensitivity of exconjugant clones to kanamycin. Kanamycin-sensitive exconjugants were screened for the mutant with colony PCR using altgenoF and altgenoR primers designed outside of the deletion flanks with the following PCR reaction mix: 10 µl of GoTaq Green master mix (Promega), 0.5 µl of each primer at 10 µM concentration, 3 µl of colony DNA template and 6 µl of sterile Milli-Q water to make a total of 20 µl single reaction. To prepare the colony DNA template for PCR, a sterile pipette tip was used to scrap kanamycin-sensitive exconjugants and suspended in 100 µl of sterile Milli-Q water. The suspension was denaturated at 95^0^C for 10 minutes. The denatured mix was centrifuged for 2 minutes. The supernatant was then used as a DNA template for PCR reaction. The PCR conditions used were: 95^0^C for 5 minutes followed by 35 cycles of 95^0^C at 20 s, 60^0^C at 30 s, 72^0^C for 1 minute followed by a final extension at 72^0^C for 5 minutes. Amplicon was expected only from the TTG mutants. The PCR amplicon was visualized in 1.5% agarose gel stained with SYBR Safe DNA gel stain (Thermo Fisher Scientific). The PCR product of the selected deletion mutant was purified using Monarch PCR and DNA cleanup kit (NEB) and sent for sequencing at Eurofins Genomics LLC (Louisville, KY, USA). The sequenced reads were aligned with the extended TTG cluster region of strain 20GA0385 to confirm the deletion mutant.

### Construction of TTG complementation plasmid

A broad host range pBBR1 derived plasmid pBBR1MCS-2 (GenBank: U23751.1) was used for building the TTG complementation plasmid. Primer pair altcomplngibF2 and altcomplngibR2 was designed to amplify the TTG gene cluster and 489 nucleotide region upstream of *altB* and 405 bp region downstream of *altI* gene region. A 35 bp Gibson overhang upstream and downstream of unique XbaI restriction site in the multiple cloning site (MCS) of the plasmid pBBR1MCS-2 was added to 5’ region and 3’ region of altcomplngibF2 and altcomplngibR2 primers respectively. The gene region was amplified from the *B. gladioli* strain FDAARGOS_389 strain using Q5 High-Fidelity DNA polymerase PCR (New England Biolabs) following 20 µl final volume and reaction components mentioned in the manufacturer’s protocol (https://www.neb.com/en-us/protocols/2013/12/13/pcr-using-q5-high-fidelity-dna-polymerase-m0491). The PCR conditions used were: 98^0^C for 30 s followed by 30 cycles of 98^0^C for 10s, 59^0^C for 30 s, 72^0^C for 4 minutes, and final extension for 2 minutes. The PCR amplicon was visualized in 1.5% agarose gel and the band obtained in the expected region was excised and purified using a Monarch NEB Gel extraction kit. The PCR product from the repeated reaction was purified using Monarch NEB PCR cleanup kit and purified product was digested with AvrII and XbaI restriction enzyme. The visualized digest pattern in the gel was compared with the simulated *in silico* pattern to confirm the TTG gene cluster-specific PCR product. The pBBR1MCS-2 plasmid was linearized with XbaI restriction enzyme and the gel product was excised and purified as described above. Gel-purified TTG PCR product and XbaI linearized plasmid were mixed in a reaction with NEB Hifi DNA assembly polymerase master mix following the manufacturer’s protocol to perform the ligation reaction. The ligation mix was then transformed into chemically competent DH5α cells and selected on an LB plate amended with kanamycin and Xgal. The clone with insert was expected to be white on Xgal plates. White clones obtained on the plates were screened for the correct insert using genotyping primer pair altcomplnF designed upstream of the attB2 site and M13R49 primer binding region in the plasmid backbone. A product size of 544 bp was expected for the correct insert. The PCR conditions used were: 95^0^C for 5 minutes followed by 35 cycles of 95^0^C for 20 s, 55^0^C for 30 s, 72^0^C for 45 s, and a final extension for 5 minutes. Similarly, a second PCR primer set altgenocomplnR was designed downstream of the attB1 site and used with M13F43 primer in the plasmid backbone to genotype the TTG insert. The expected product size was 514 bp. PCR was performed in 20 µl reaction using GoTaq green master mix with reaction mix followed as described before. The PCR conditions used were: 95^0^C for 5 minutes followed by 35 cycles of 95^0^C for 20 s, 62^0^C for 30 s, 72^0^C for 45 s, and a final extension for 5 minutes. The insert was also confirmed using BsrGI restriction digest and whole plasmid sequencing at Plasmidsaurus (Plasmidsaurus, Eugene, OR).

For the construction of TTG Type B complementation plasmid, a primer pair with 35 bp overhang from upstream and downstream sequence region of unique XbaI restriction site was designed targeting 6842 bp of the TTG cluster gene region in the strain BC93_12. Colony PCR was performed in a 20 µl reaction using the Q5 polymerase protocol as described in the manufacturer’s protocol. The following PCR conditions were used: 98^0^C for 30 s followed by 30 cycles of 98^0^C for 10s, 61^0^C for 30 s, 72^0^C for 210 s, and final extension for 2 minutes. The gel purified PCR product was ligated to XbaI linearized pBBR1MCS-2 plasmid using NEB Hifi DNA assembly reaction and the reaction mixture was transformed to chemically competent DH5α cells and selected on LB plates amended with kanamycin and Xgal. Colonies appearing white on the Xgal kanamycin plates were screened for the insert using primer pair typeBgenocomplnR designed upstream of the attB2 sequence and standard primer M13F43 in the plasmid backbone. The expected product size was 627 bp. PCR conditions followed were the same as described above for primer pair altgenocomplnR and M13F43. A plasmid clone with the expected amplicon size was sent for sequencing at Plasmidsaurus.

### Construction of TTG expression clone in TTG negative natural variant strains background

The sequenced confirmed Type A and TTG Type B gene clusters in pBBR1MCS-2 plasmid were transformed to electrocompetent RHO5 cells and selected on LB plate amended with DAP and kanamycin following incubation at 37^0^C overnight. No DNA control was also included. The pBBR1MCS-2 Empty Vector (EV) was also transformed into electrocompetent RHO5 cells and selected following the same procedure. Single transformant Type A, Type B, and EV colonies growing on LB DAP kanamycin plate were inoculated to start an overnight culture and conjugated with representative TTG negative *Burkholderia* natural variant strains using biparental mating. *B. cepacia* natural variant BC83_1 conjugants were selected on LB plate amended with 1000 µg of kanamycin per mililitre, *B. orbicola* natural variant LMG 30279 TTG conjugants were selected on LB plate amended with 200 µg of kanamycin per mililitre whereas *B. gladioli* natural variant BG92_3 TTG conjugants and engineered 20GA0385 TTG mutants were selected on LB amended with rifampicin and 50 µg of kanamycin. Each of the TTG negative natural variant strains and engineered TTG mutants were conjugated with pBBR1MCS-2 plasmid harboring both Type A and TTG Type B clusters. The insert was confirmed using the genotyping primers described in the previous section following the same procedure.

### Preparation of allicin stock solution

The allicin stock preparation procedure was followed as described by (Stice et al., 2020) with slight modifications. Briefly, 15 µl of diallyl disulfide 96% (Carbosynth), 25 µl of glacial acetic acid (Sigma Aldrich), and 15 µl of 30% H_2_0_2_ were mixed in a 200 ul PCR tube. The tube was sealed with parafilm, attached to a 500 ml beaker, and agitated at a 28^0^C shaker for 6 h. The reaction mix was then suspended in 1 ml of methanol. The methanol allicin mix was used as a synthesized allicin stock for the ZOI assay.

### Zone of inhibition assay

The ZOI assay was performed to test the quantitative/qualitative differences in resistance between the strains harboring endogenous or plasmid-based TTG clusters relative to the TTG-negative engineered strains or natural variant strains. The LB overnight cultures (O/N) (∼24 hr) amended with appropriate antibiotics were started from a single colony of the representative strains. Polystyrene petri plates (100 mm x 15 mm) with 20 mL of LB agar and appropriate antibiotics as needed were spread with 300 µl of bacterial suspension. All natural variant strains were plated on LB media. *B. gladioli* BG92_3 strain was plated on LB amended with rifampicin and its TTG plasmid derivatives were plated on LB amended with kanamycin and rifampicin. *B. cepacia* TTG negative variant was plated on LB and its TTG plasmid derivatives were plated on LB amended with 1000 µg/ml kanamycin. *B. orbicola* strain LMG 30279 TTG plasmid derivatives were plated on LB amended with 200 µg/ml kanamycin plate and its WT was plated on LB. The WT *B. gladioli* strain 20GA0385 and its engineered TTG mutant derivative were plated on LB and the Type A and TTG Type B plasmid complement derivatives were plated on LB amended with rifampicin and kanamycin for the ZOI assay. Three plate replicates were used per treatment. A circular well was poked at the center of the plate using the back end of sterile 10 µl pipette tips and 50 µl of the synthesized allicin stock was added to the wells. Plates were incubated at 28°C for 24 h. The ZOI area was measured using ImageJ software (Abràmoff et al., 2004). The experiment was repeated at least 3 times. The statistical difference in the ZOI area among different natural variants within a species was determined using one-way Analysis of Variance (ANOVA) and Tukey’s Honestly Significant Difference (HSD) test using agricolae library and the box plot was generated using ggplot2 function in RStudio v 2023.9.0.

### Preparation of onion juice

Yellow onions were purchased from the grocery store and the crude juice was extracted following the procedure described by Stice et al., 2020. Briefly, a consumer-grade juicer (Breville Juice Fountain Elite) was used to crush the onion bulb which yielded 200-300 ml of crude onion extract. The extract was centrifuged (Sorvall RC5B Plus, Marshall Scientific, Hampton, NH) in a 250 ml centrifuge bottle (14000 g, 2 h, 4°C). The upper phase liquid was sterilized using a Nalgene disposable 0.2 micron vacuum filter sterilization unit. The prepared juice was aliquoted and stored at -20^0^C for less than 1 week for experimental use.

### Preparation of bacterial inoculum

Bacterial inocula for the ZOI assays, onion foliar assays, RSN assays, and foliar and scale time course experiments were prepared following the same procedure for all tested *Burkholderia* strains. The test strains were streaked on an LB plate amended with appropriate antibiotics and incubated at 28°C for 24 h. A day after, colony growth from the lawn was suspended in 300 µl of sterile Milli-Q water and incubated for 24 h overnight. A day after, colony growth from the lawn was scooped and suspended in 1 ml of ddH_2_0 /MgCl_2_ and standardized to OD_600_ = 0.7 (∼2.4 x 10^8 CFU/ml, *B. gladioli* 20GA0385). The standardized suspension was used for inoculation.

### Onion filtered extract growth assay

The growth assay experiment was performed in 100-well honeycomb plates using the Bioscreen C system (Lab Systems, Helsinki, Finland). The standardized suspension volume of 40 µl was added to 360 µl of half-strength filtered onion extract (180 µl of onion juice diluted with 180 µl of sterile ddH20). Each honeycomb well was loaded with 380 µl of the mix and each treatment had six well replicates. The bioscreen experiment was run for 48 h with shaking at 28°C and the absorbance values were recorded every 30 minutes. The experiment was repeated three times for each tested species. Statistical analysis of OD_600_ values at each time point for different treatments was performed using the pairwise t-test function in RStudio 2023.09.0. The average OD_600_ reading of water-onion extract negative control treatment for each time point was subtracted from the OD_600_ reading values obtained for each well for all the treatments at the corresponding time point.

### Onion foliar/seedling necrosis assay

Onion seedlings (*Allium cepa* L. cv Texas grano 1015 Y supersweet onions) of 8-12 weeks were used for the foliar assay. Inoculum for strains 20GA0385 pBBR1MCS-2 EV, 20GA0385 ΔTTG pBBR1MCS-2 EV, 20GA0385 ΔTTG pBBRR1MCS-2:: TTG Type A, BG92-3 and its Type A and TTG Type B plasmid derivatives were prepared as previously described. Onion seedlings were trimmed to keep the oldest blade intact and approximately the midpoint of the blade was poked on one side with a sterile 20 µl pipette tip to create a wound. The normalized bacterial suspension of 10 µl prepared with 0.25 mM MgCl_2_ was deposited into the wounded tissue. Negative controls were inoculated with sterile 0.25 mM MgCl_2_. Maximum lesion length was measured 3 days post inoculation (dpi). Each treatment had six biological repeats and the experiment was repeated at least three times.

For quantification of bacterial population in the infected blade tissue, a section of the infected tissue measuring 0.5 cm above and below the point of inoculation was cut and resuspended in 200 µl of Milli-Q H_2_0 in a 2-ml SARSTEDT microtube (SARSTEDT AG & Co., Numbrecht, Germany). The tissue was manually crushed using a sterile blue pestle and then ground with a SpeedMill PlUS homogenizer (AnalytiK Jena) two times for 1 minute each. To facilitate the maceration, three 3-mm zirconia beads (Glen Mills grinding media) and one 4.5 mm bead (store-bought) were added to the tissue and water mix. Serial dilutions were performed using 10 µl of the ground tissue and diluents were plated on LB plates amended with rifampicin and kanamycin. The number of colony-forming units (CFUs) was back-calculated to determine the bacterial population levels in the infected tissue.

The time course foliar assay was performed using 4-6 months old onion plants (cv. Century) grown in the greenhouse. *B. gladioli* strain 20GA0385 and its TTG mutant derivative were used for the study. The onion blades were trimmed to keep the oldest two leaves intact that were used for inoculation. The midpoint of the blade (measured from the tip to the base of the blade) was marked with a sharpie and a sterile 20 µl pipette tip was used to poke a hole on one side of the leaf to create wounding. Bacterial inoculum concentration of OD_600_ = 0.7, ∼2.4 x 10^8^ colony forming units (CFU)/ml for *B. gladioli* 20GA0385 WT and the TTG mutant, was used. Normalized bacterial suspension (10 µl) was deposited to the wounding site. Three plants were inoculated per treatment (six blades total). A total of 36 onion plants (72 blades in total) were inoculated and sampled daily from 0 to 5 dpi. Day 0 samples were processed for bacterial quantification 6 hours post-inoculation.

The bacterial population in the infected tissue was quantified by sampling the tissue samples 0.5 cm above and 0.5 cm below the point of inoculation. The tissue was weighed in 200 µl of sterile Milli-Q H_2_0 in a 2-ml SARSTEDT microtube. The tube was filled with 3-mm zirconia beads for maceration with a GenoGrinder and an additional 4.5 mm bead was added for grinding with the SpeedMill PlUS homogenizer. For grinding with GenoGrinder, 30 seconds was used and the SpeedMill homogenizer was used for maceration two runs of 1 minute each. The resulting macerate was serially diluted to 10^8^ in 0.25 mM MgCl_2_ in 96-well styrene plates (20 µl, 180 µl). The dilutions were plated on LB square plates amended with rifampicin and cfu per mg was back-calculated for each treatment. The cfu/mg value for two inoculated leaves from the same plant was averaged to get a single cfu/mg value per biological replicate per treatment. The log folded cfu/mg values of three biological replicates per treatment were plotted against days post-inoculation. The statistical difference in bacteria population levels each day between the two treatments was analyzed using a pairwise t-test function in RStudio. The experiment was repeated two times.

### RSN assay

The RSN assay was set up following the procedure described in (Stice et al., 2018), and (Shin et al., 2023) with slight modifications. Red onion bulbs were purchased from a grocery store and sliced into 3-5 x 3-5 cm-sized scales. The scales were surface sterilized in a 3% sodium hypochlorite solution for 2 minutes and rinsed in deionized water six times. After drying on a sterile paper towel for a few minutes, the scales were placed on an ethanol wiped pipette tip rack. Each scale was wounded using a sterile 20 µl pipette tip, and then inoculated with 10 µl of a standardized bacterial suspension. Sterile 0.25 mM MgCl_2_ was inoculated as a negative control. The scales were placed on a flat potting tray (27 x 52 cm) lined with two layers of paper towel that had been moistened with 50 ml of deionized water. Another flat was placed on top of the tray to maintain humidity and the entire setup was incubated at room temperature for 72 h. Six scales were inoculated per treatment and the experiment was repeated three times.

The scale necrosis area after 72 h was measured using ImageJ. For quantification of bacteria in the necrotic onion tissue, approximately 0.5 x 0.5 cm area around the point of inoculation was excised with a sterile scalpel, suspended, and weighed in a 1.5 ml microcentrifuge tube with 200 µl of sterile water after 4- and 72-hour post-inoculation. The tissue was crushed manually with the end of a sterile wooden cocktail stick. The resulting macerate was diluted ten-fold in a dilution series with sterile 0.25 mM MgCl_2_ in 96-well styrene plates (20 µl, 180 µl). The diluents (10 µl) were plated on LB amended with rifampicin and kanamycin as appropriate and incubated at 28°C for 36 h. Colonies were counted and cfu per milligram was back-calculated for each sampled scale. Three scales treated with the negative control were processed 4 h post-inoculation and three were processed 72 h post-inoculation. For the rest of the treatments, six inoculated scales were processed. Statistical analysis of the differences between species-specific treatments was performed using the pairwise t-test function in RStudio.

The inoculum preparation, assay setup, and quantification of bacteria in infected onion tissues for the RSN time course experiment were conducted as described in the previous section. The *B. gladioli* strain 20GA0385 pBBR1MCS-2 EV and its TTG mutant derivative were used for the experiment. The bacterial suspension was normalized to OD_600_ value of 0.7 followed by 100-fold dilution. Each scale was inoculated with 10 µl of diluted suspension. A total of 72 individual scales were inoculated and six scales per treatment were sampled daily from day 0 to day 5 post-inoculation. Subsequently, a dilution series ranging from 10^-1^ to 10^-3^ of the TTG mutant treatment was plated on an LB square plate amended with rifampicin and kanamycin. The statistical analysis of the difference in log_10_ fold cfu per milligram bacterial recovery between the two treatments at each time point was performed using a pairwise t-test function in RStudio.

## Acknowledgements

We acknowledge George Sundin for graciously proving the *B. orbicola* strain AU 1054, Gi Yoon Shin and all the Kvitko, Dutta, and Yang lab members for their helpful comments and suggestions during the preparation of this manuscript.

---

**Table S1:**
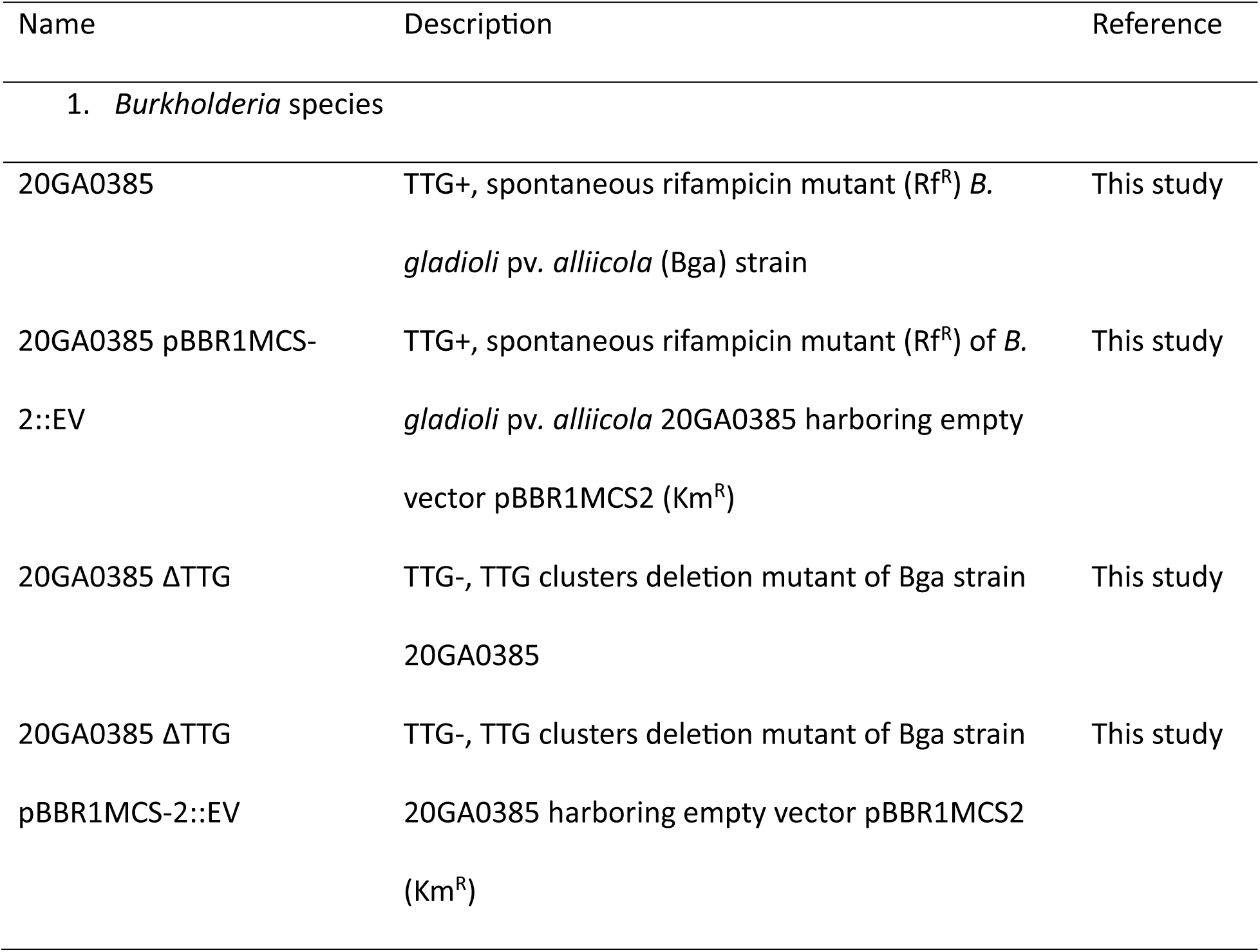
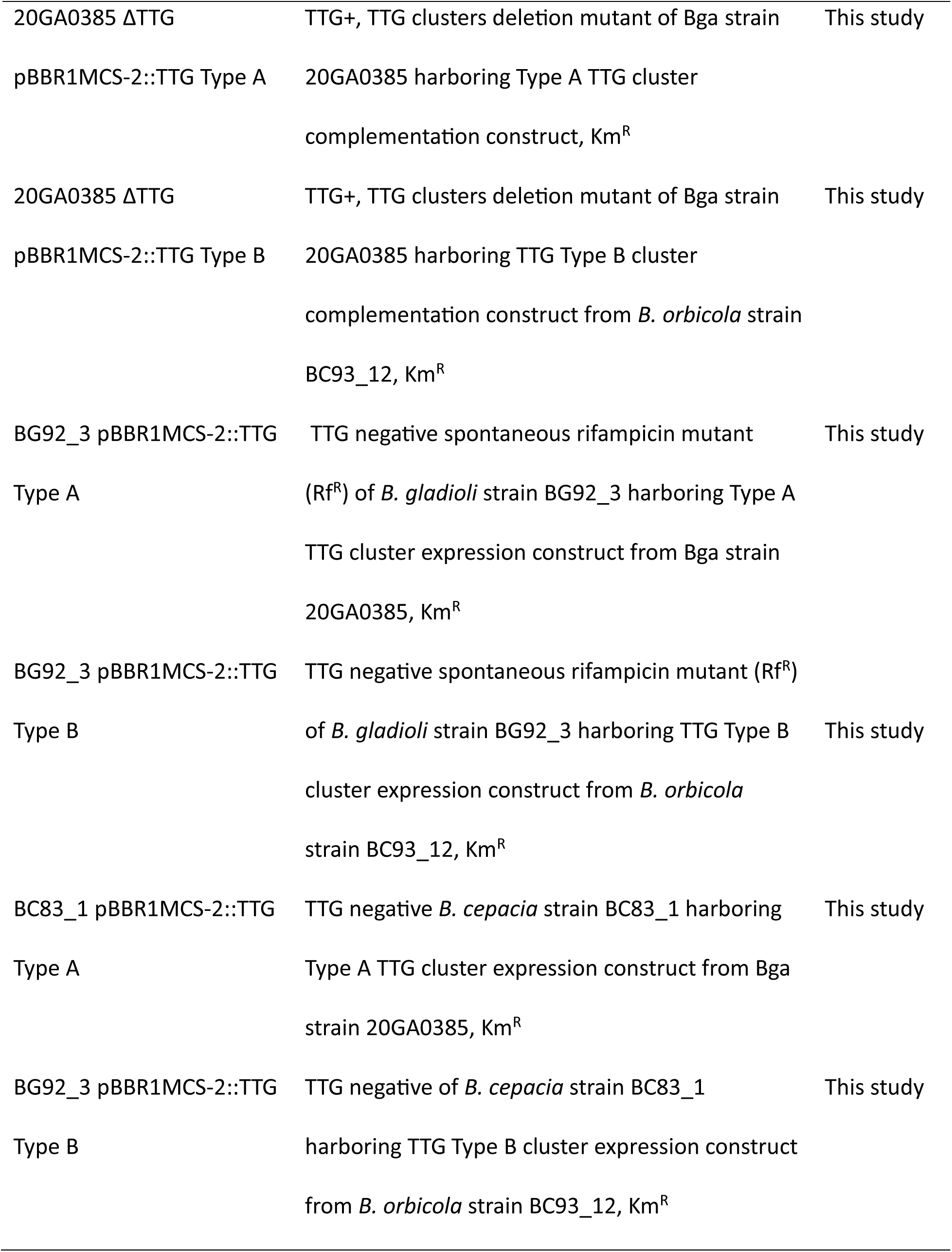
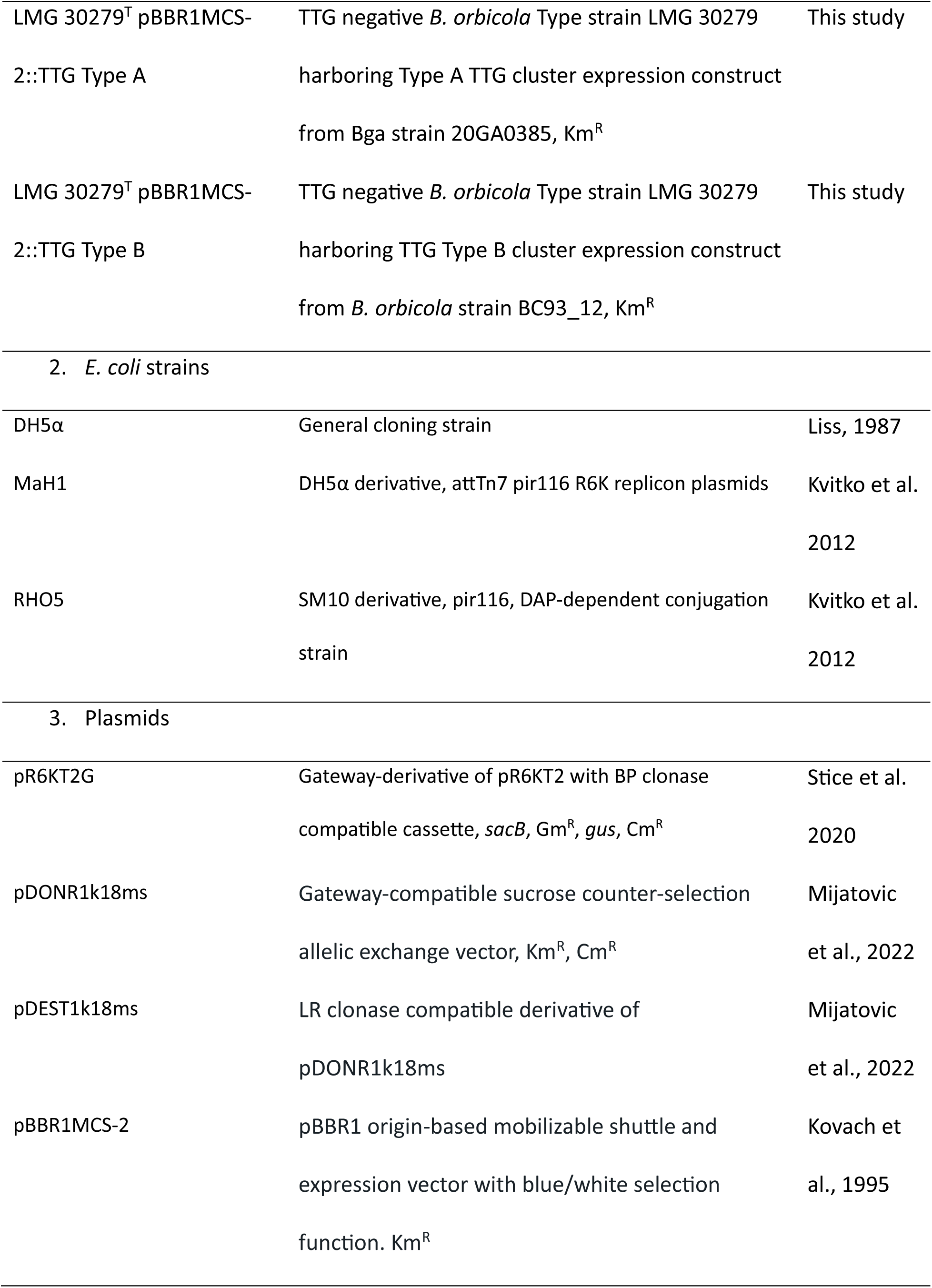
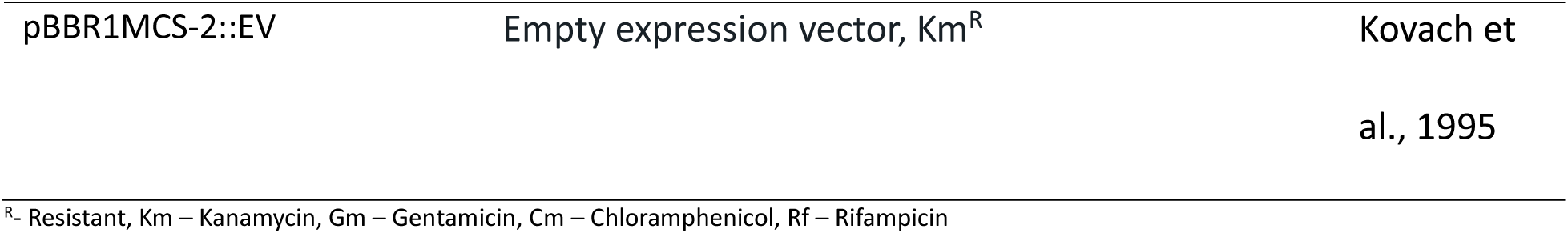
Bacterial strains and plasmids used in cloning and mutagenesis.

**Table S2:**
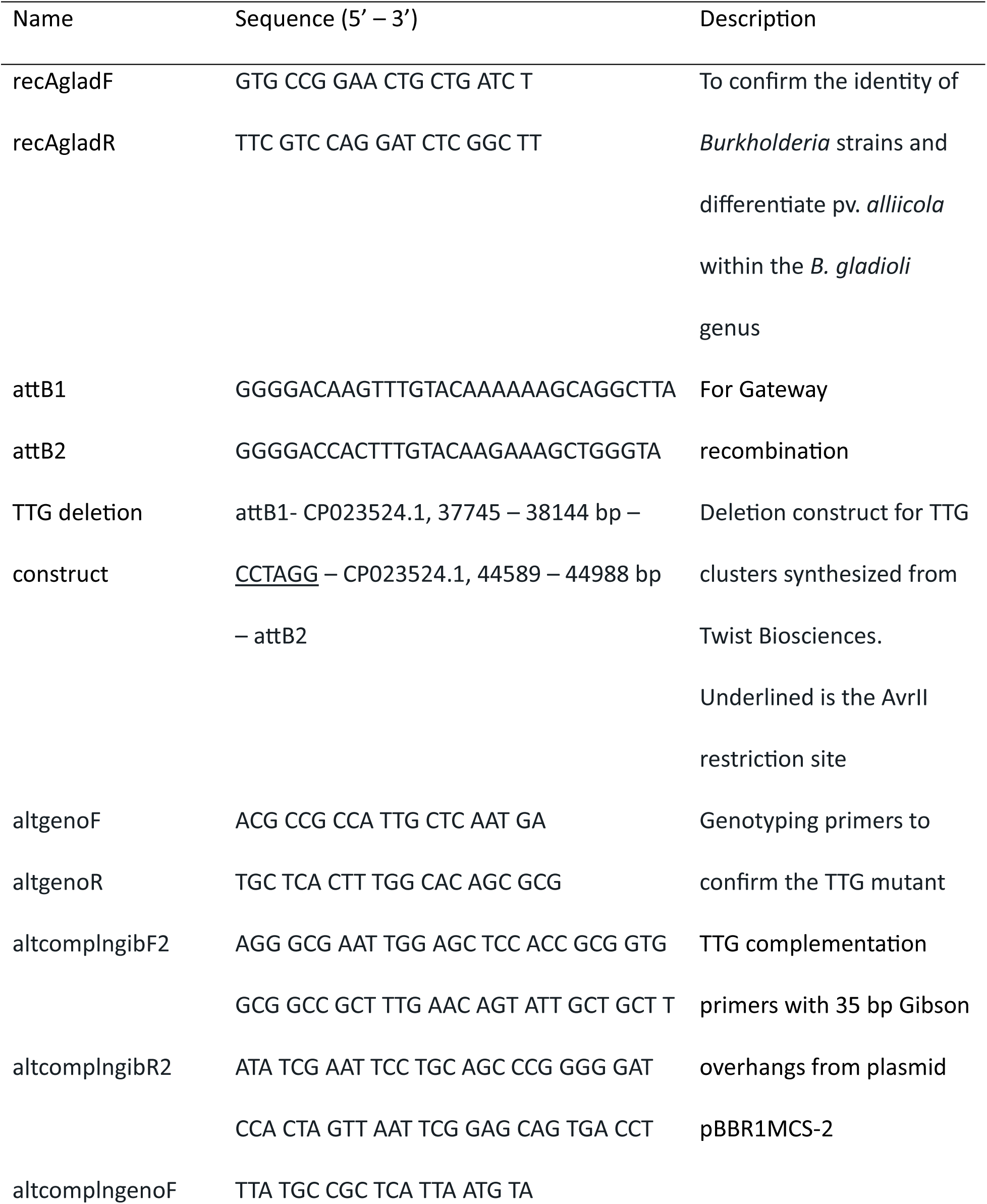
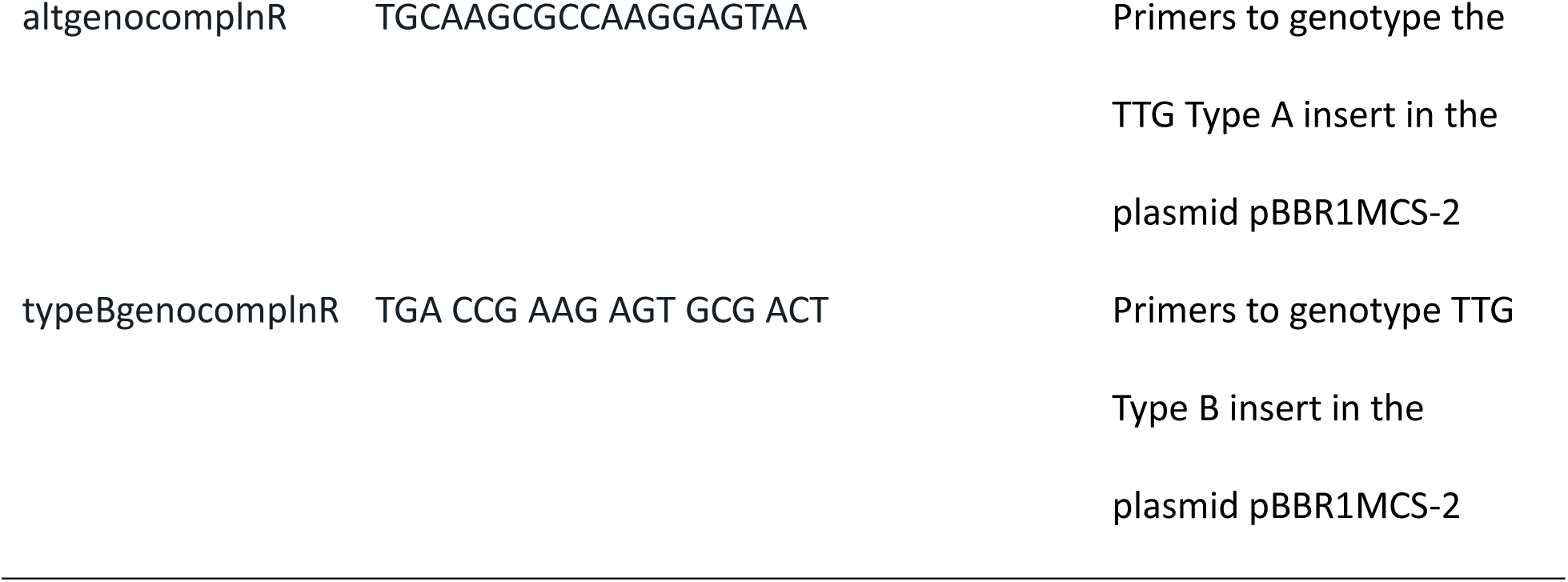
Primers and dsDNA gene blocks used in the study.

**Table S3:**
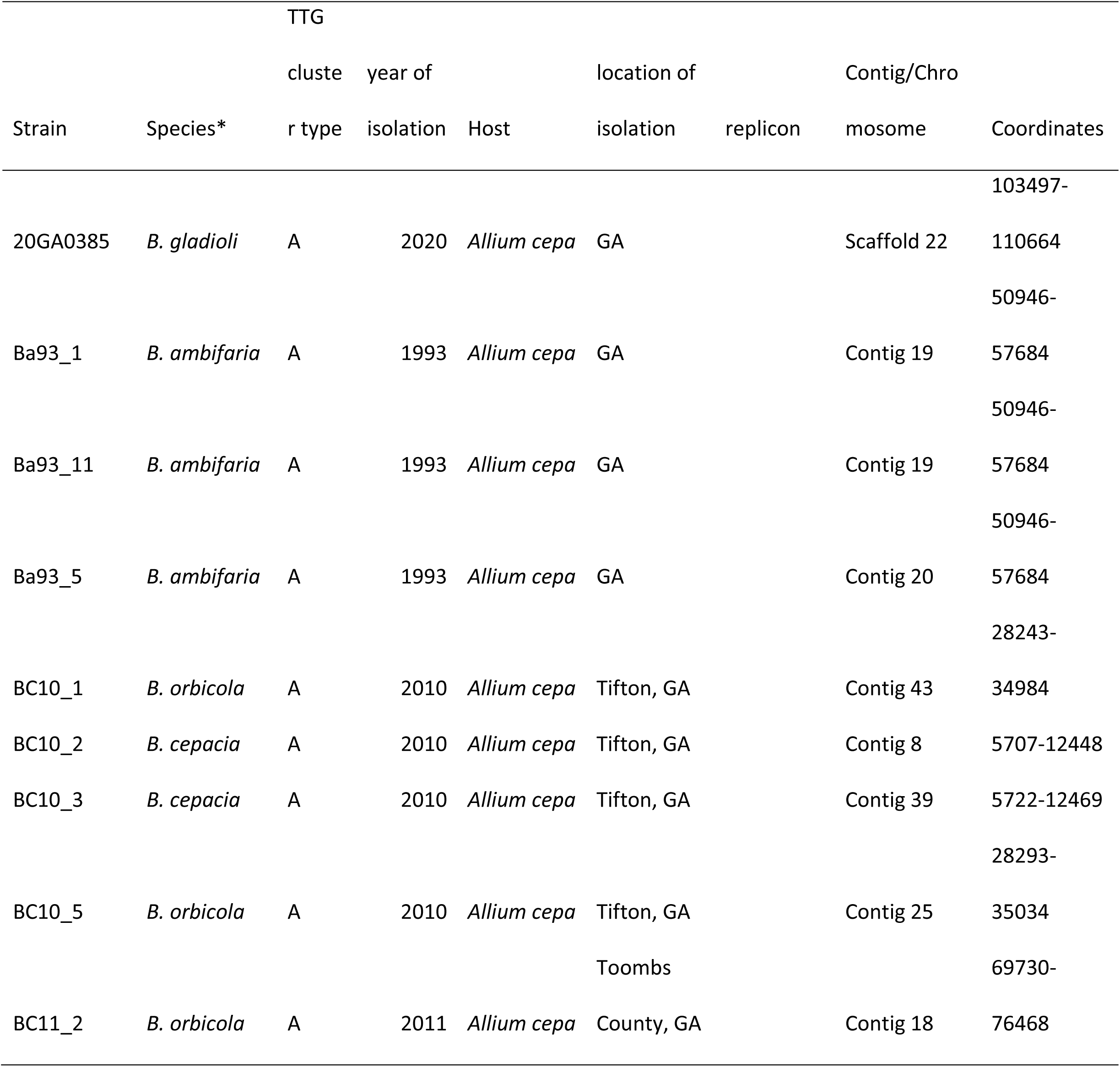
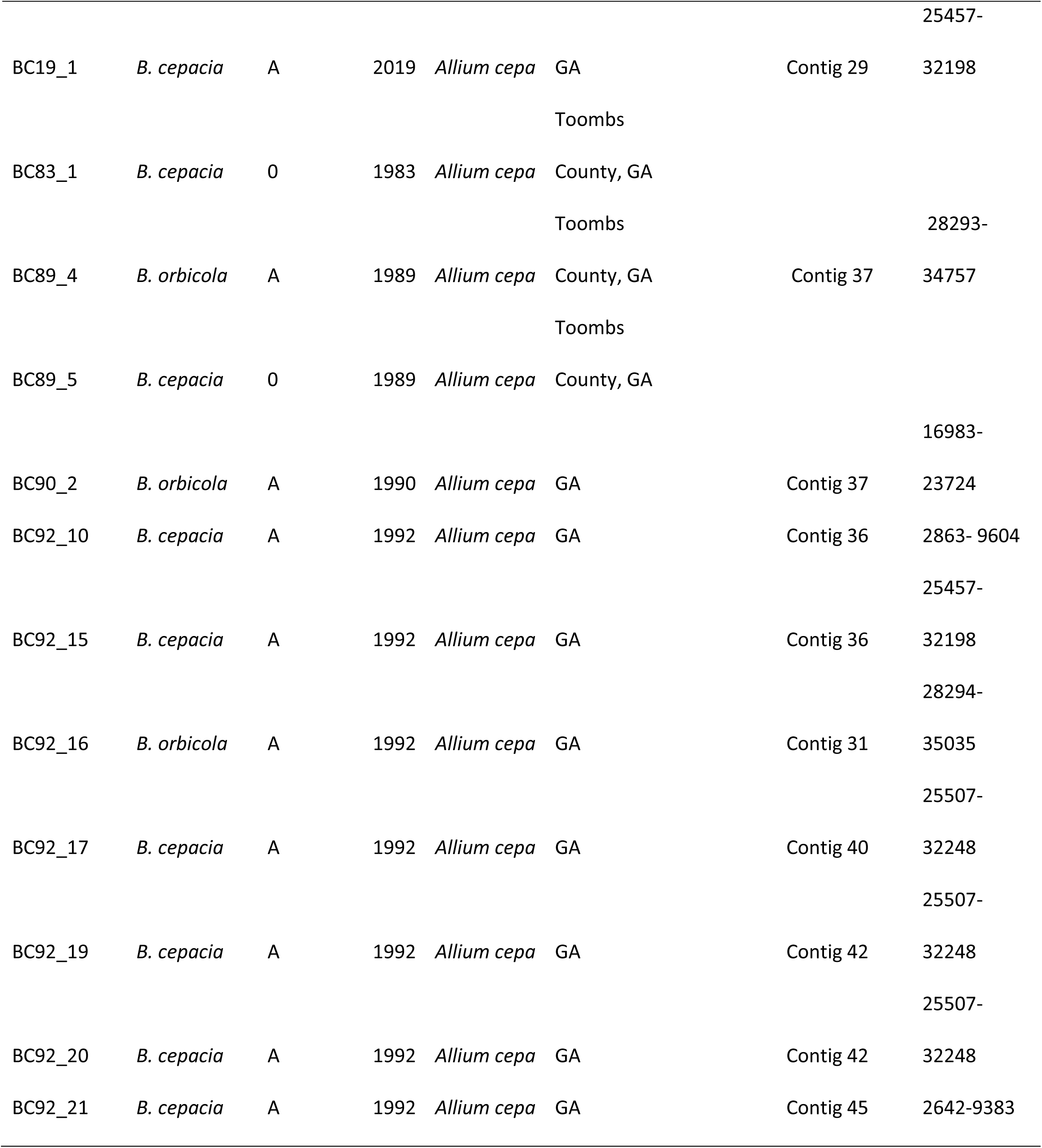
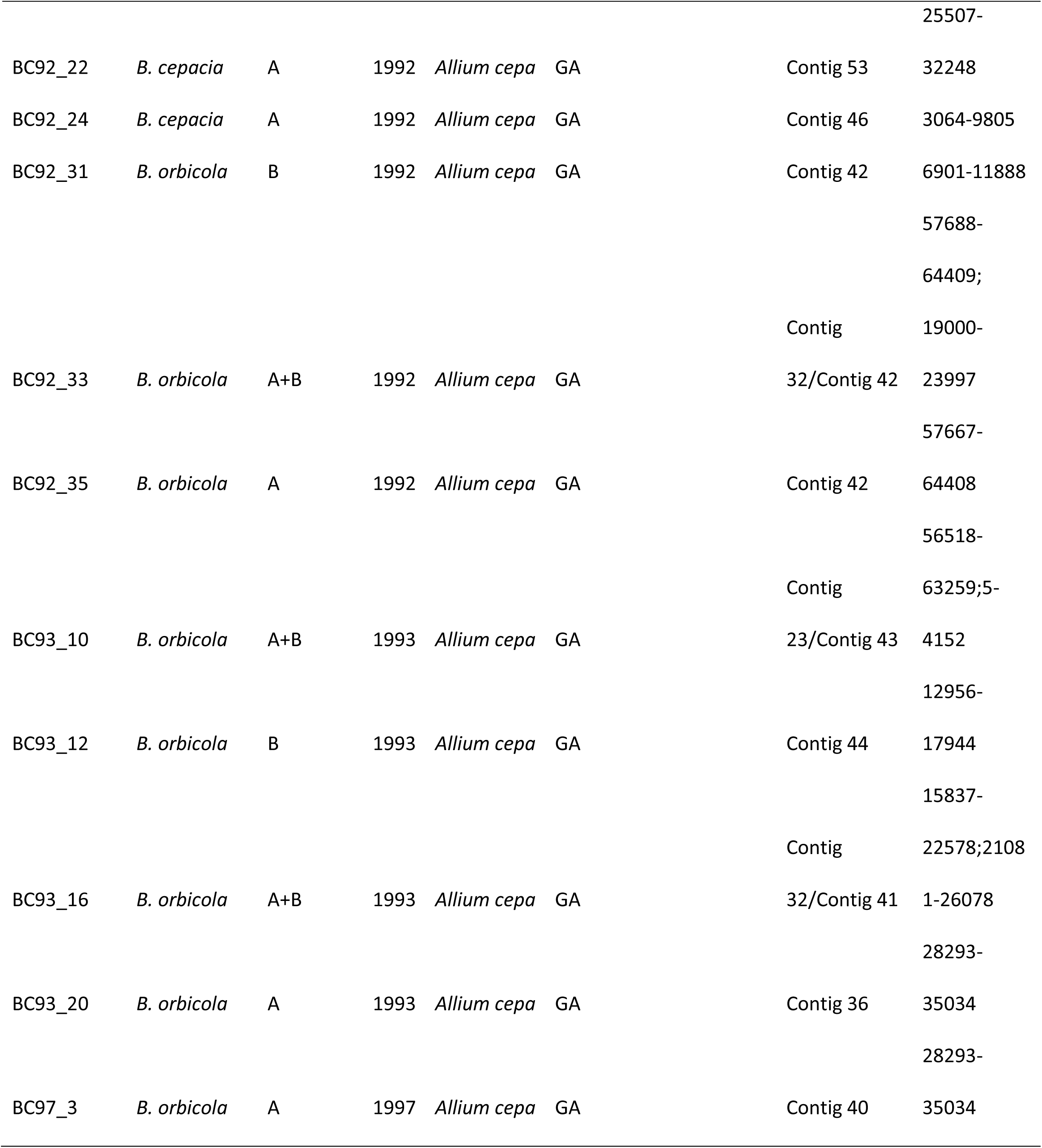
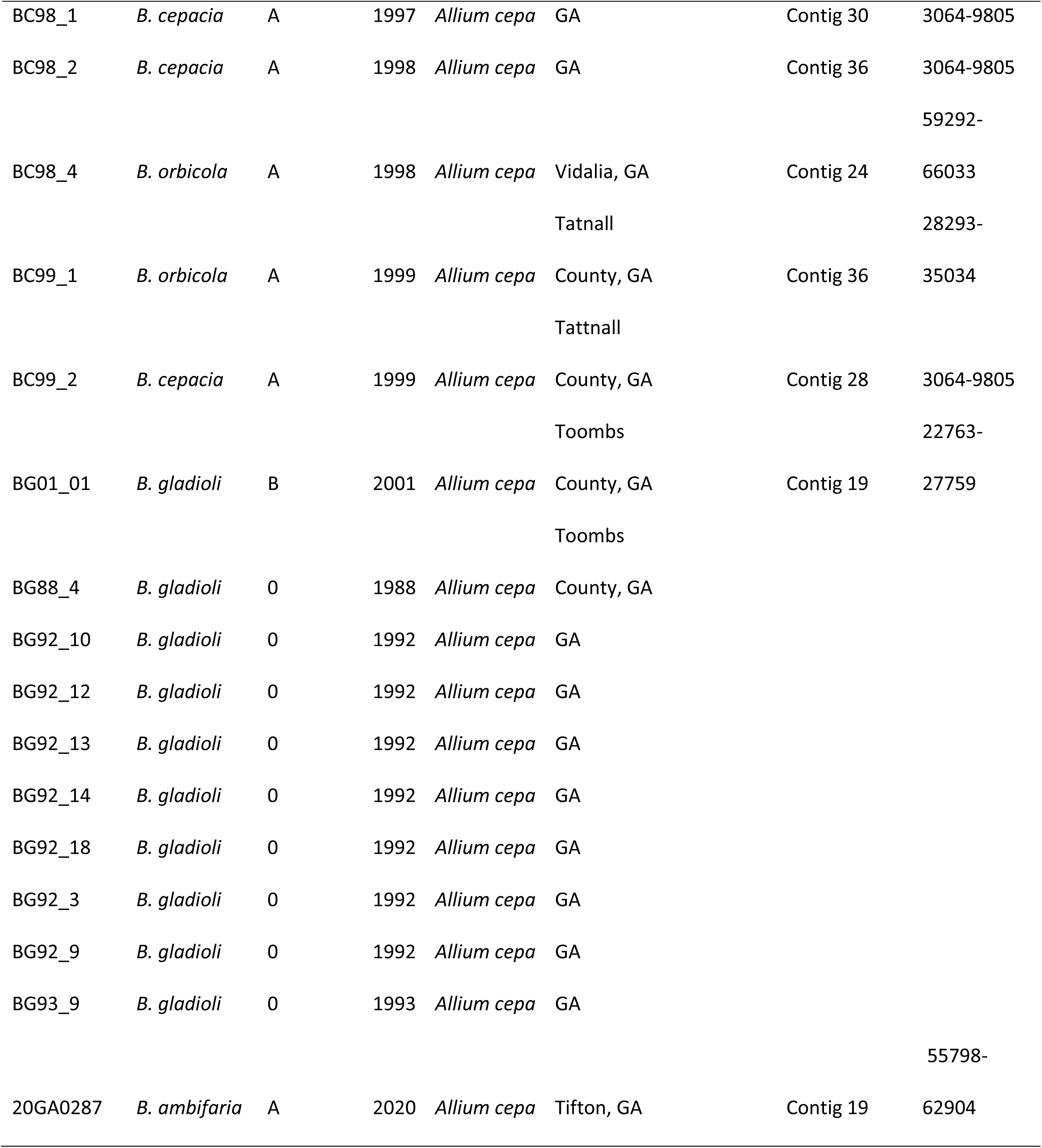
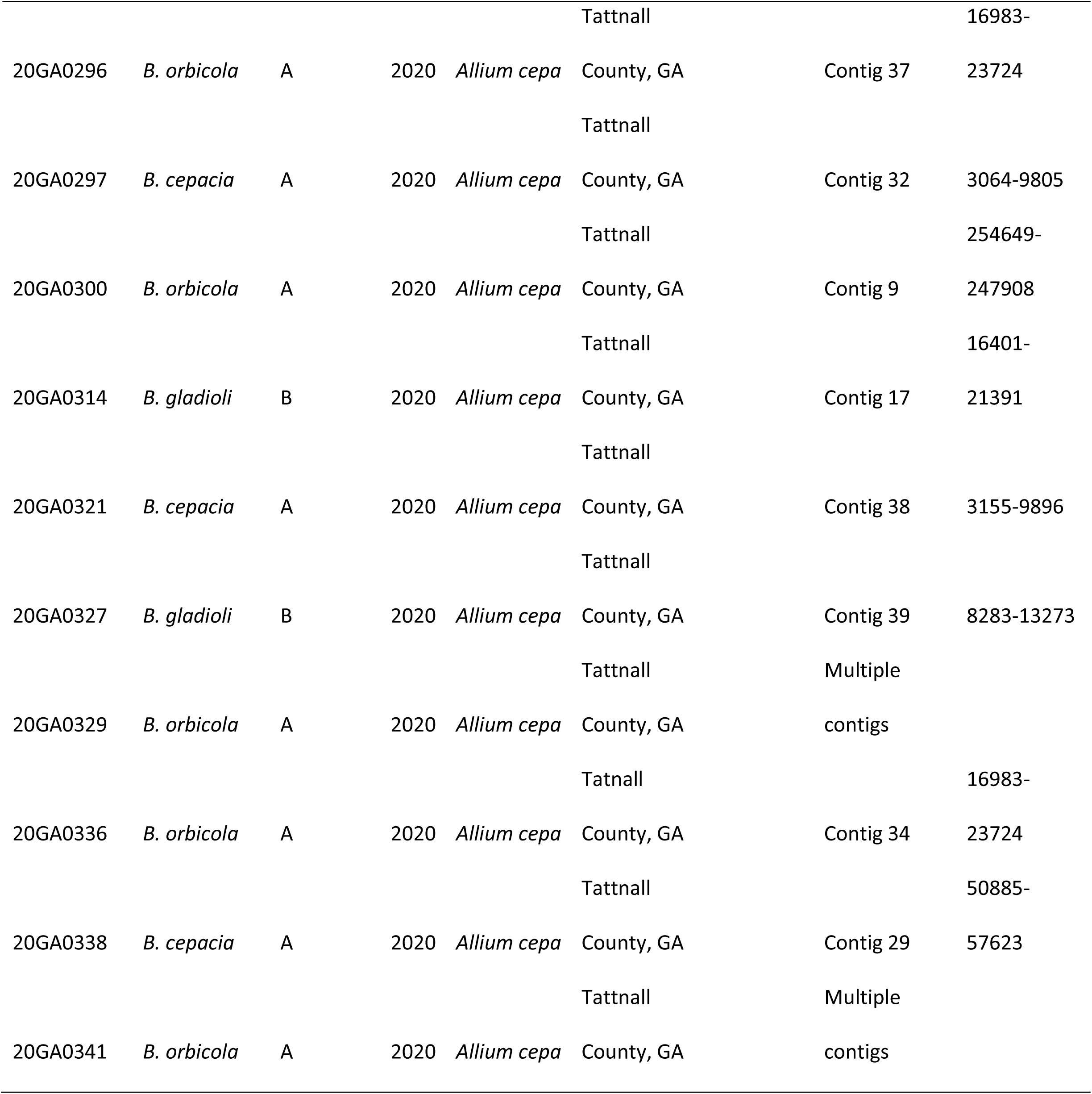
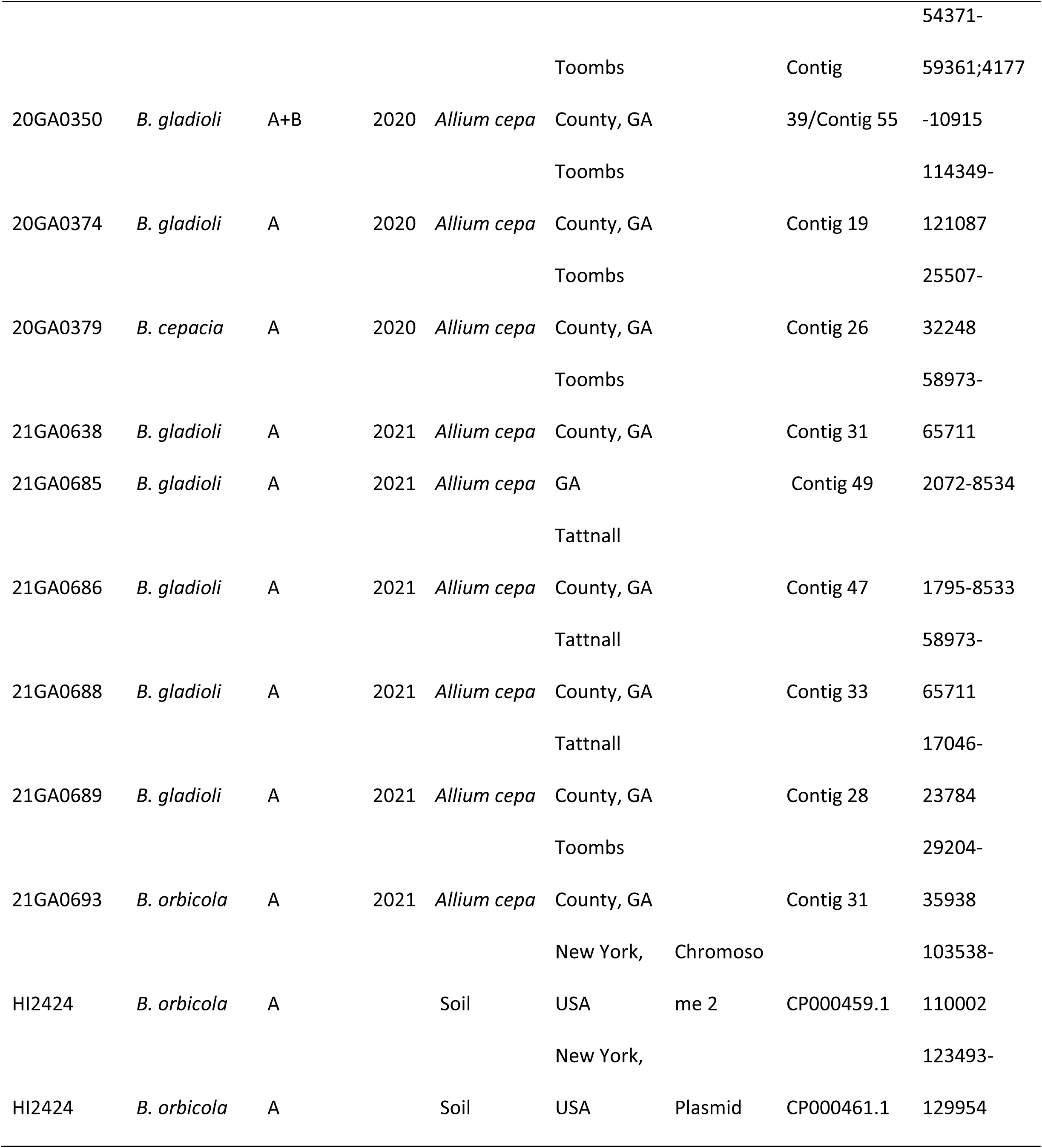
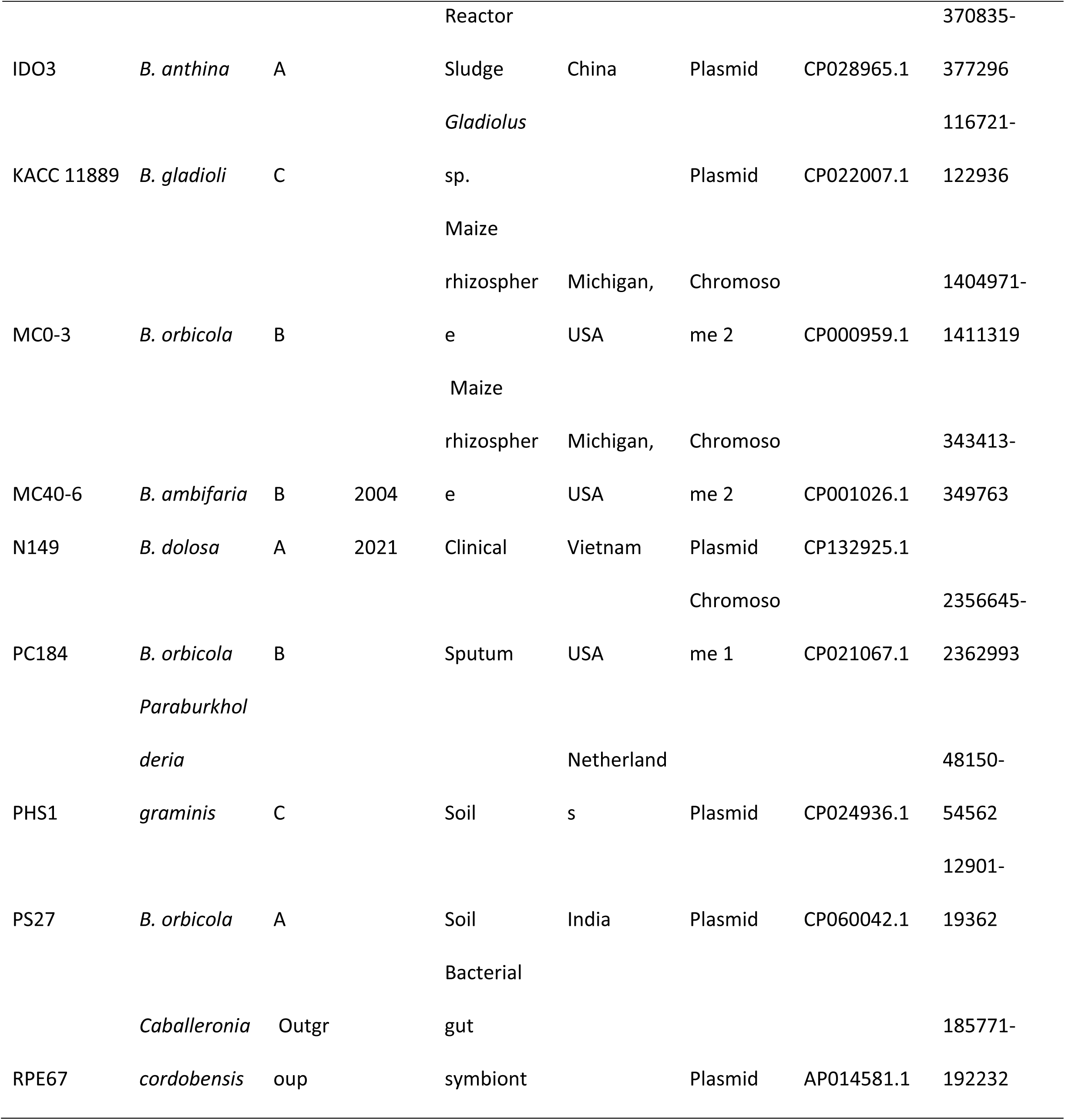
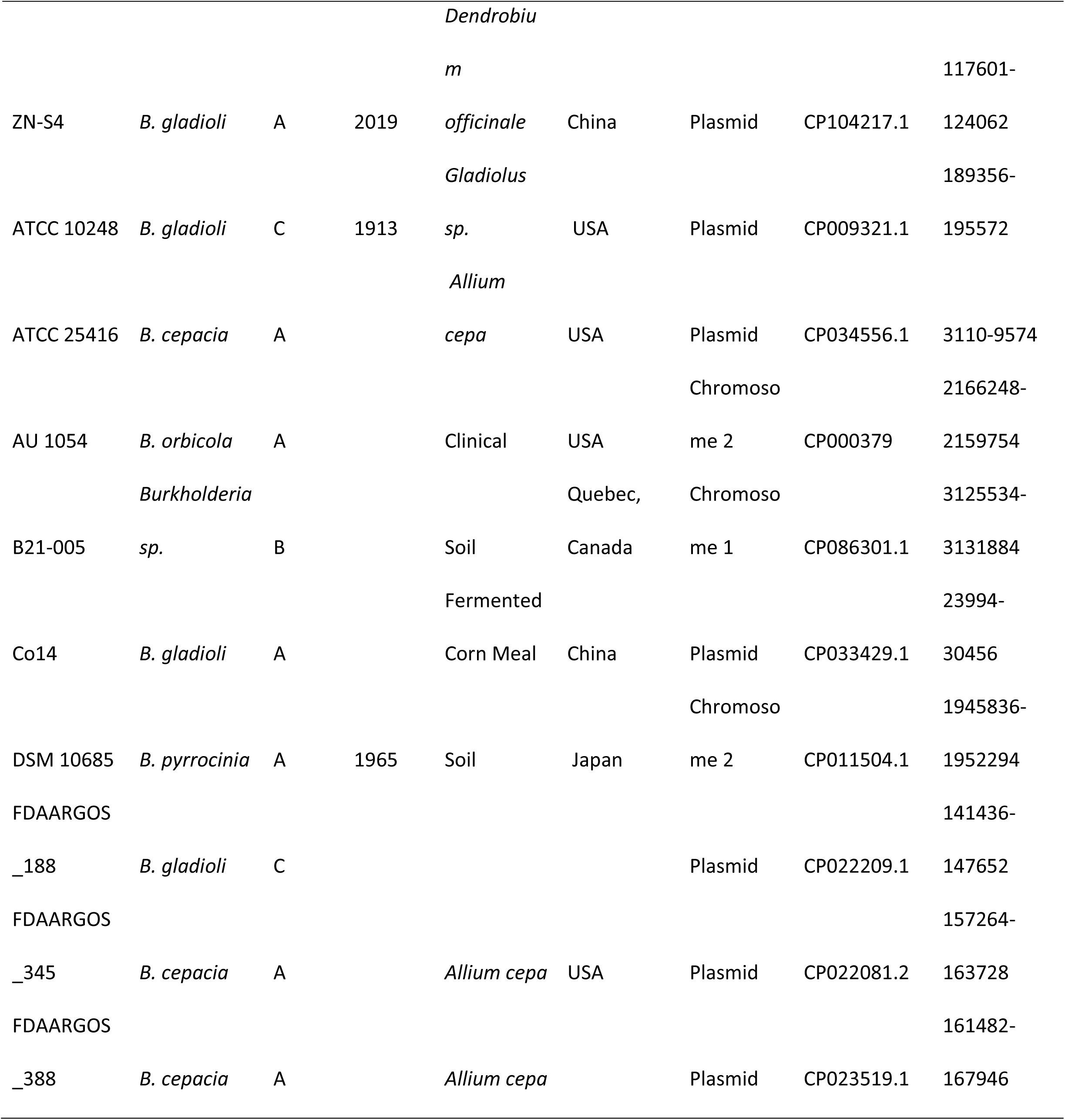
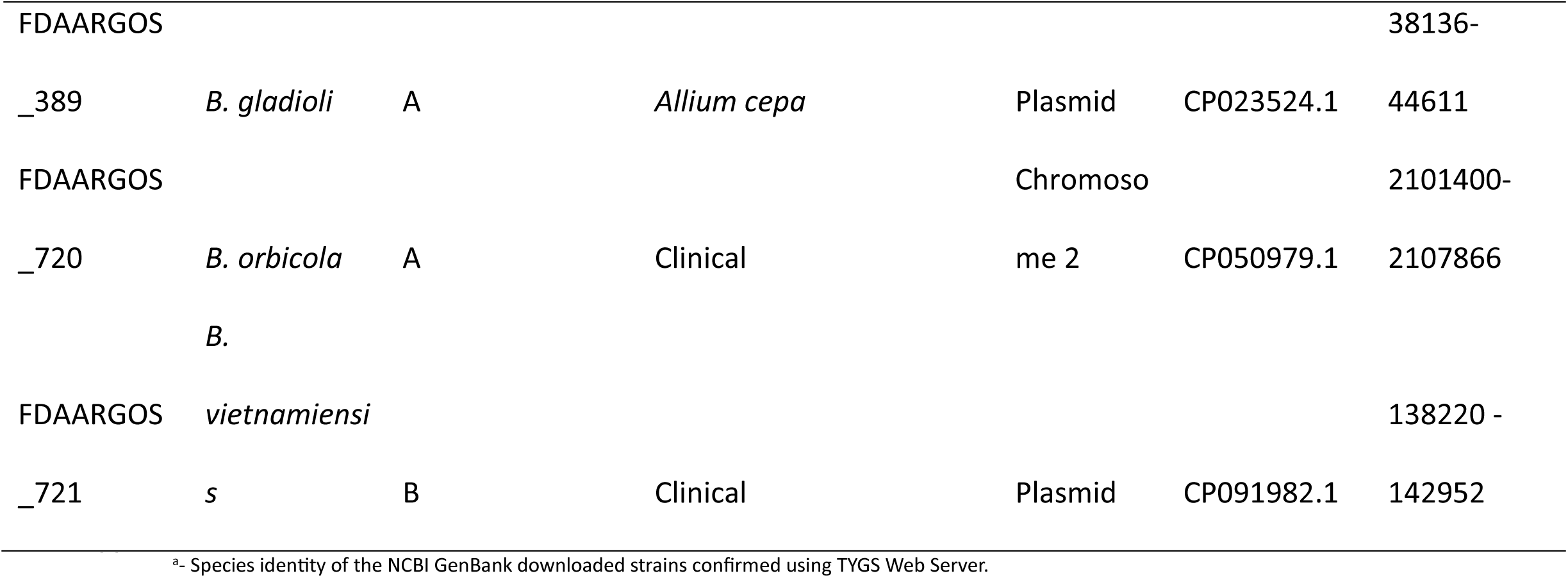
Description of the whole genome sequenced and NCBI GenBank extracted strains used for the phylogenetic analysis.

**Table S4:** Level of significance between OD_600_ values of different treatments at different time points across the three experimental repeats of onion juice assay. The average value of six data points of Onion Juice water negative control at each time point of each experimental repeat is subtracted from every single well of other treatments at the same corresponding time point. Total number of data points n = 18 per treatment. Analysis was done using a pairwise t-test function in RStudio.

| Hours | 0.5 | 4 | 8 | 12 | 16 | 20 | 24 | 28 | 32 | 36 | 40 | 44 | 48 |
| --- | --- | --- | --- | --- | --- | --- | --- | --- | --- | --- | --- | --- | --- |
| *B. gladioli* 20GA0385 ΔTTG Vs |  |  |  |  |  |  |  |  |  |  |  |  |  |
| WT | ns | ns | ns | ns | ns | ns | ** | *** | *** | *** | *** | *** | *** |
| ΔTTG p<sup>TTGA</sup> | ns | ns | ns | ns | ns | *** | *** | *** | *** | *** | *** | *** | *** |
| *B. cepacia* BC83_1 (TTG-) vs |  |  |  |  |  |  |  |  |  |  |  |  |  |
| p<sup>TTGA</sup> | ns | *** | *** | *** | *** | *** | *** | *** | *** | *** | *** | *** | *** |
| p<sup>TTGB</sup> | ns | *** | *** | *** | *** | *** | *** | *** | *** | *** | *** | *** | *** |
| *B. cepacia* BC83_1 p<sup>TTGA</sup> vs |  |  |  |  |  |  |  |  |  |  |  |  |  |
| p<sup>TTGB</sup> | ns | * | ns | ns | ** | ns | ns | ns | ns | ns | ns | ns | ns |
| *B. orbicola* LMG 30279 (TTG-) vs |  |  |  |  |  |  |  |  |  |  |  |  |  |
| p<sup>TTGA</sup> | ns | ns | ns | ns | ns | *** | ** | *** | *** | *** | *** | *** | *** |
| p<sup>TTGB</sup> | ns | ns | ns | *** | *** | *** | *** | *** | *** | *** | *** | *** | *** |
| *B. orbicola* LMG 30279 p<sup>TTGA</sup> vs |  |  |  |  |  |  |  |  |  |  |  |  |  |
| p<sup>TTGB</sup> | ns | ns | ns | ns | ** | *** | *** | *** | *** | *** | *** | *** | *** |

ns – non-significant, level of significance: 0 '\*\*\*' 0.001 '\*\*' 0.01 '\*' 0.05

**Supplementary Figure S1:** Maximum likelihood phylogenetic tree of 727-bp partial *recA* gene differentiates *Burkholderia gladioli* pv. *alliicola* group from other pathovars in *B. gladioli. B. gladioli* 20GA0385 and NCBI GenBank extracted FDAARGOS_389 test strains for identity confirmation are represented by starburst symbol. Information about the strain and their pathovar designation is retrieved from Jones et al., 2021. The letter/number in parentheses represents the clade of the reference strain. The black circle represents bootstrap support of the branching on a scale of 0 to 1. T denotes a type strain. The tree is rooted in the branch of *B. cepacia* complex representative strains.

