## Supplementary figures and images for "Thiosulfinate tolerance gene clusters are common features of *Burkholderia* onion pathogens"

### Figure S1

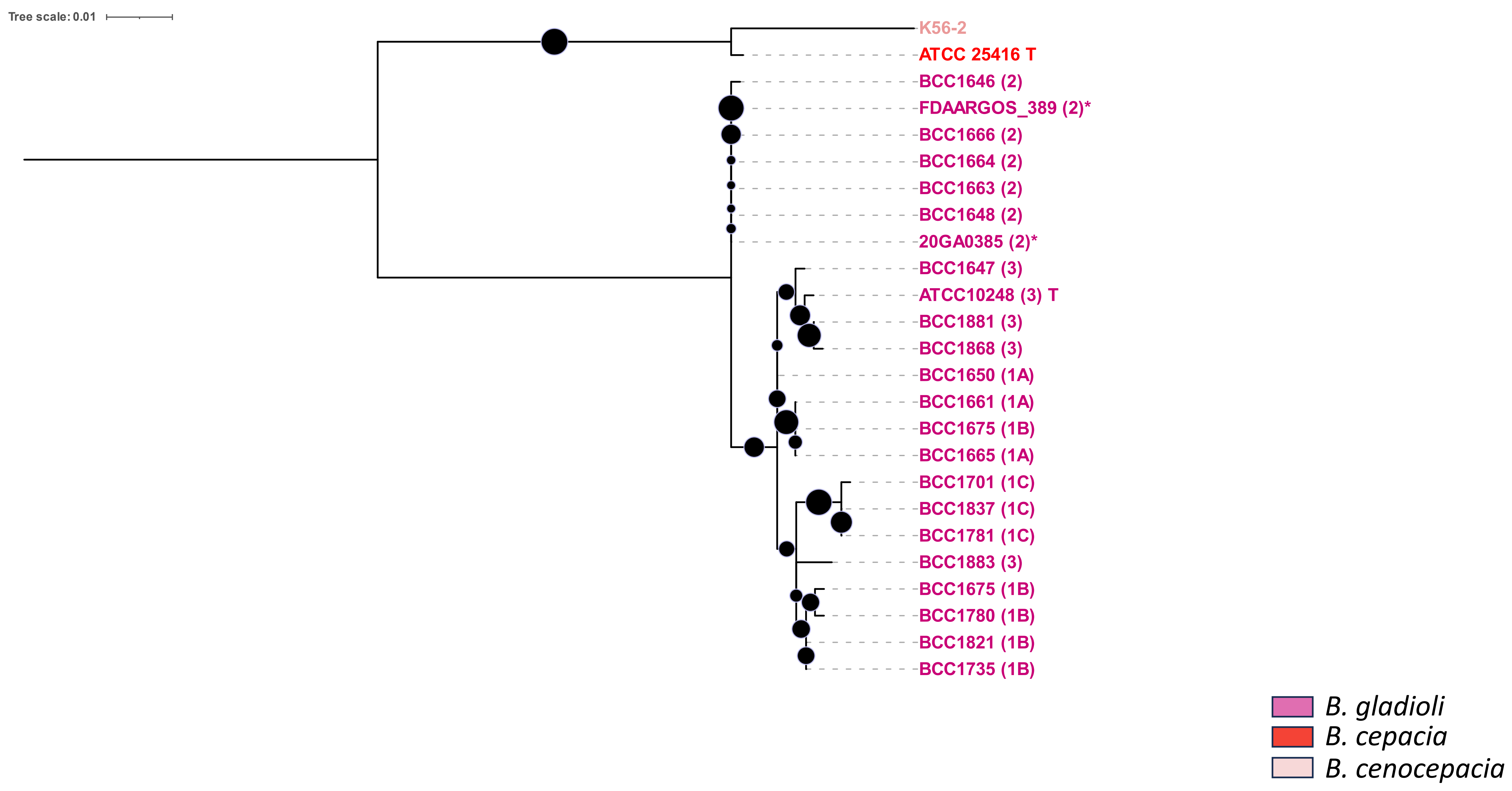
